# Rabbit Development as a Model for Single Cell Comparative Genomics

**DOI:** 10.1101/2022.10.06.510971

**Authors:** Mai-Linh N. Ton, Daniel Keitley, Bart Theeuwes, Carolina Guibentif, Jonas Ahnfelt-Rønne, Thomas Kjærgaard Andreassen, Fernando J. Calero-Nieto, Ivan Imaz-Rosshandler, Blanca Pijuan-Sala, Jennifer Nichols, Èlia Benito-Gutiérrez, John C. Marioni, Berthold Göttgens

**Affiliations:** Department of Haematology, University of Cambridge, Cambridge, UK; Wellcome-Medical Research Council Cambridge Stem Cell Institute, University of Cambridge, Cambridge, UK; Department of Zoology, University of Cambridge, Cambridge, UK; Inst. Biomedicine, Dept. Microbiology and Immunology, Sahlgrenska Center for Cancer Research, University of Gothenburg, Gothenburg, Sweden; Department of Pathology & Imaging, Novo Nordisk, Måløv, Denmark; Medical Research Council Laboratory of Molecular Biology, Cambridge, UK; Genome Biology Unit, European Molecular Biology Laboratory (EMBL), Heidelberg, Germany; MRC Human Genetics Unit, Institute of Genetics and Cancer, University of Edinburgh, Edinburgh, UK; Wellcome Sanger Institute, Wellcome Genome Campus, Cambridge, UK; European Molecular Biology Laboratory, European Bioinformatics Institute, Cambridge, UK; Cancer Research UK Cambridge Institute, University of Cambridge Cambridge, UK

## Abstract

Biomedical research relies heavily on the use of model organisms to gain insight into human health and development. Traditionally, the mouse has been the favored vertebrate model, due to its experimental and genetic tractability. Non-rodent embryological studies however highlight that many aspects of early mouse development, including the egg-cylinder topology of the embryo and its method of implantation, diverge from other mammals, thus complicating inferences about human development. In this study, we constructed a morphological and molecular atlas of rabbit development, which like the human embryo, develops as a flat-bilaminar disc. We report transcriptional and chromatin accessibility profiles of almost 180,000 single cells and high-resolution histology sections from embryos spanning gastrulation, implantation, amniogenesis, and early organogenesis. Using a novel computational pipeline, we compare the transcriptional landscape of rabbit and mouse at the scale of the entire organism, revealing that extra-embryonic tissues, as well as gut and PGC cell types, are highly divergent between species. Focusing on these extra-embryonic tissues, which are highly accessible in the rabbit, we characterize the gene regulatory programs underlying trophoblast differentiation and identify novel signaling interactions involving the yolk sac mesothelium during hematopoiesis. Finally, we demonstrate how the combination of both rabbit and mouse atlases can be leveraged to extract new biological insights from sparse macaque and human data. The datasets and analysis pipelines reported here set a framework for a broader cross-species approach to decipher early mammalian development, and are readily adaptable to deploy single cell comparative genomics more broadly across biomedical research.

## INTRODUCTION

Due to the ethical and technical challenges of experimenting with human and non-human primate embryos, our understanding of human development is largely influenced by studies on the mouse. The mouse has dominated as a mammalian model for over 50 years for its ease of maintenance and genetic manipulation, short generation times and large litter sizes. However, despite the utility of the mouse model in recapitulating many aspects of human biology, it has long been recognized that there is significant variability in early development across mammals and particularly between rodent and non-rodent species. A prominent example is the egg-cylinder shape of the mouse embryo, which substantially deviates from the flat-disc morphology of most other amniotes, including humans, and even other rodents ^1^. As a result of the cup-shape morphology, the germ layers have an inverted organization compared to flat-disc species, with early endoderm on the outside of the embryo. These cells subsequently internalize through a rodent-specific turning process to establish the phylotypic vertebrate body plan ^2^. The mouse embryo also diverges in other respects, such as the topology of its extra-embryonic tissues ^3^ and method of implantation ^4^.

Discordance between human and animal models is one of the primary reasons pre-clinical pharmaceuticals fail efficacy or safety testing^5^. Thalidomide, for example, causes fetal malformations in rabbits, non-human primates, and humans, but not in mice ^6–8^. Differences in developmental toxicity between mice and non-rodents are indeed widely recognized, and current regulatory guidelines require that embryo–fetal developmental toxicity (EFDT) testing is conducted on both a rodent and non-rodent species^9^.

Despite this, and although rabbits, dogs, sheep, pigs, and non-human primates are all used in biomedical research, the availability of deep molecular data in mammals is largely limited to the mouse. Previous molecular studies have characterized, at single-cell resolution, gastrulation and subsequent organogenesis in the mouse, enabling delineation of specific lineage commitment events ^10, 11^ and facilitating an understanding of the role of the epigenome in cell fate commitment ^12, 13^. However, with advances in genome engineering and molecular profiling technologies^14^, more organisms have become accessible to functional experimentation and disease modeling. This presents an opportunity to gain a comparative understanding of species at the molecular and cellular level, which will be critical to determine optimal model systems, improve the translation of animal studies, and gain deeper insights into early mammalian development more broadly.

As an alternative to the mouse, the European rabbit, *Oryctolagus cuniculus*, offers many advantages for studying early mammalian development. Like mice, rabbits have short reproductive cycles (31 days), large litter sizes, and are well-established as a laboratory animal in pharmacological, reproductive, and developmental research ^15–17^ with rabbits being the most commonly used non-rodent species in developmental toxicity studies. Moreover, relative to mice, the rabbit embryo is highly accessible at later stages of development due to its large size and late implantation ^15^. The embryo implants superficially, rather than intrusively embedding into the uterine lining, making it tractable to dissect and obtain embryos. Rabbits are also phylogenetically well-positioned, sharing a more recent common ancestor to both rodents and primates than other mammalian model organisms, such as the cow or pig ^18^. Finally, compared with mice, phylogenetic models predict a smaller branch length from the Glire ancestor, suggesting that the rabbit genome is also more representative of the ancestral condition ^19, 20^. Taken together, the more conserved genetic scaffold, the morphological similarity of early development, and the ready availability of experimental material, makes studying developmental processes in the rabbit appealing, especially as a model of early human development. However, at present, a deep molecular characterisation of early development in the rabbit, and a thorough molecular comparison of rabbit to humans and mouse is lacking.

In this paper, we characterize rabbit development, using high-resolution whole-embryo histology, single-cell transcriptomics and single-cell chromatin accessibility profiling, at three gestational days (GDs) 7-9 that capture implantation, amniogenesis, and gastrulation. Using a novel neighborhood-based approach, we place the rabbit transcriptional landscape in context with the mouse, identifying conserved and divergent cell states, cell types and developmental trajectories across species. We also investigate the molecular programmes underlying trophoblast differentiation and yolk sac hematopoiesis in the rabbit and use the fine-grained cell type annotations of both rabbit and mouse atlases to obtain a more detailed view into the cell type diversity of sparse human and macaque data. Our transcriptional and chromatin accessibility atlases can be explored via the interactive website available through https://marionilab.github.io/RabbitGastrulation2022/.

## RESULTS

### A time-resolved single cell RNA-seq atlas of rabbit gastrulation and early organogenesis

Gestational Days (GD) 7, 8 and 9 of New Zealand White Rabbit development encompass implantation, gastrulation and early organogenesis (Figure 1A-C). Individual embryos across each of these stages were processed to obtain an anatomical and morphological view of rabbit embryogenesis via high-resolution histology imaging on transverse and sagittal sections (Figure 1C; Methods). Up until the blastocyst stage, rabbit and mouse embryos share a largely similar structure with an outer layer of trophectoderm surrounding epiblast and hypoblast cell layers^21^. However, in contrast to the mouse embryo, which implants around day 4.5 ^22^, the rabbit blastocyst fills with fluid and expands drastically prior to implantation. The polar trophoblast, known as Rauber’s layer, deteriorates in rabbits, so the epiblast is directly exposed to the maternal environment and sits on top of this large bag of fluid (Figure 1C, S1A)^23^. These patterns can be clearly observed in our histological reference set: at GD7 the expanded blastocyst and the first signs of implantation are observed on the anti-mesometrial side of the embryo (Figure 1A,C, S1A,D)^24^. Additionally, the primitive streak can clearly be seen and the flat-disk shaped gastrula is similar to the shape of a CS7 human gastrula ^25^. By GD8, the first somites can be seen (Figure 1A) and the amnion regrows from the surrounding trophoblast (Figure 1C, S1B,F,H). Contrasting with mouse development, this is the stage of development at which implantation completes on the mesometrial side. Finally, by GD9, the earliest structures corresponding to more mature organs can be seen, such as the optic vesicle, heart, neural tube, and allantois (Figure 1A,1C, S1F).

**Figure 1:**
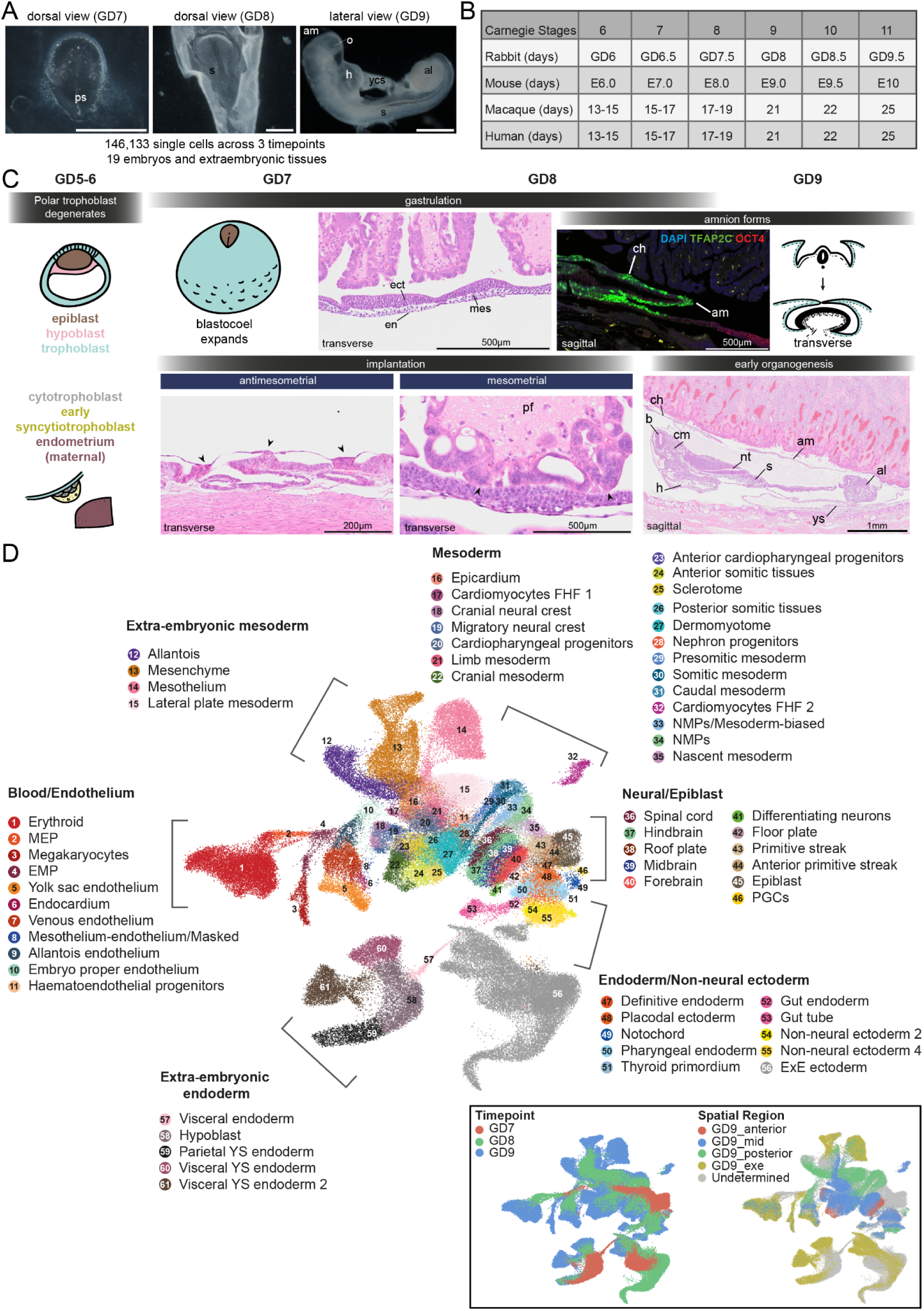
Rabbit Development at a Glance. **a)** images of dissected embryos at gestational day (GD) 7, 8, and 9. Scale bar = 1mm. ps = primitive streak, s = somites, am = amnion, o = optic vesicle, h = heart, ycs = yolk connecting stalk, al = allantois **b)** Relative Carnegie stages of rabbit, mouse, macaque, and human and corresponding day of development. **c)** Timeline with schematics and images of key developmental events that occur between GD6-9 in the rabbit. Implantation and gastrulation occur simultaneously in the rabbit from GD7-8, with anti-mesometrial implantation occurring first, marked by the adherence of trophoblastic knobs to the uterine lining (indicated with arrows). This is shortly followed by mesometrial implantation at GD8. Image shows implantation sites (arrows) in the mesometrial placental fold. By GD9, early organs are formed such as the fetal heart, amnion, allantois, and brain. Histology images representative of each process are shown as well as RNAscope images to show the amniochorionic fold at GD8 with RNA probes for TFAP2C, OCT4 and DAPI staining to show the nucleus. ect = ectoderm, en = endoderm, mes = mesoderm, ch = chorion, b = brain, cm = cranial mesoderm, nt = neural tube. **d**) UMAP of 146,133 cells captured in a transcriptional atlas, labeled by annotated cell type. box: the same UMAP labeled by (left) time point and (right) anatomical region, based on the microdissection of GD9 embryos. MEP = megakaryocyte–erythroid progenitors; EMP = erythro-myeloid progenitors; FHF = first heart field; NMPs = neuromesodermal progenitors; PGCs = primordial germ cells; ExE = extraembryonic; YS = yolk sac.

Having generated this anatomical reference, we next looked to molecularly profile cells of the rabbit embryo across these developmental stages. 6 individual embryos for GD7, 3 individual embryos and 2 pools of 3 embryos each for GD8, and 4 embryos for GD9 were processed for single-cell RNA-seq using the 10X Genomics Chromium System (Methods, Figure S3A). All GD9 embryos were dissected into the embryo-proper and extraembryonic region; two of which were micro-dissected further into the anterior, mid, and posterior regions providing high-level spatial information (Figure S1C). Following quality control and initial processing, including substantial improvement of the rabbit transcriptome annotation (Methods, Figure S2, S3), we obtained high-quality transcriptome profiles for 13,674 cells at GD7, 34,686 cells at GD8 and 97,773 cells at GD9, providing deep transcriptomic sampling along this entire period of rapid embryo growth (Figure 1D, S3).

To assign each profiled cell to a specific cell type, we employed a combination of automated and manual annotation approaches. In a recent atlas of mouse gastrulation and early organogenesis, 430,339 cells spanning embryonic day (E) 6.5 to E9.5, were classified into 87 curated cell type labels ^26^. Given the overlapping developmental stages (Figure 1B), we trained a label-transfer model on the full mouse dataset and used this to predict cell type annotations for the rabbit (Methods, Figure S4A). The automatic transfer of existing annotations ensured that our rabbit cell types were defined as consistently as possible with the mouse, facilitating cross-species comparisons downstream. The preliminary annotations obtained from this automated stage were subsequently verified by manual annotation via assessment of known cell type markers and by cross-referencing with additional information such as the developmental stage, histology (Figure 1A, S1D) spatial information via embryo microdissection (Figure S1C), RNA-scope images and independent data integration (Figure S4C). Novel cell types not shared between species were also identified in this stage through examining marker genes, high-resolution in situ hybridization microscopy (Figure S4B), and thorough consultations with specialist collaborators. In total, we defined 67 cell types across a library of 146,133 cells which encompass different lineages arising from the three earliest precursors, namely the epiblast, hypoblast and trophoblast (Figure 1D).

### Chromatin and gene-regulatory dynamics capture key processes of trophoblast-syncytiotrophoblast differentiation in vivo

From the fertilized zygote, the first fate decision is between the inner cell mass (ICM) and the trophectoderm. The trophectoderm can be separated into the top, polar trophoblast layer overlying the ICM, and the bottom mural trophoblast of the blastocyst. It is noteworthy that in species such as rabbit, cat, dog, sheep, and pig, the polar trophoblast layer degenerates so that the epiblast becomes contiguous with the mural trophoblast ^23^. The remaining trophoblast cells subsequently differentiate to form the amniotic ectoderm, cytotrophoblast and syncytiotrophoblast, which mediate implantation and restructuring of the maternal environment to accommodate the developing embryo.

Despite being of vital importance for the successful development of the embryo, the gene-regulatory programmes underlying trophoblast differentiation and the establishment of the fetal-maternal interface in vivo remain poorly understood. Moreover, differences in trophoblast cell types and their key regulators between mice and humans motivate the need for additional in-vitro and in-vivo models ^27^, particularly as several pregnancy-related disorders, including preeclampsia and intrauterine growth restriction, are associated with abnormal trophoblast development ^28^. One reason why the process of implantation has been so difficult to study at the molecular level is that, in humans and mice, it takes place very early in development (prior to gastrulation), with consequent challenges in capturing enough cells for rigorous molecular profiling. Moreover, in organisms where the embryo is deeply embedded within the uterine lining (such as human and mouse), capturing trophoblast cell types without also capturing a lot of maternal material is experimentally challenging. By contrast, in the rabbit, implantation occurs alongside gastrulation at GD 7 and 8^4, 24^, when the embryo is larger, enabling the straightforward capture of large numbers of extraembryonic-ectoderm cells (Figure 1D, S1A-B).

Consistent with this, in comparison to the mouse gastrulation dataset ^10^ which only captures a subset of cells labeled as extraembryonic-ectoderm, we are able to profile a diverse set of extraembryonic ectoderm cell types (Figure 2A), encompassing the maturation of early trophoblast cells (characterized by expression of *GATA3*, *HAND1*, and *DPPA3*) ^29, 30^ into cytotrophoblast (characterized by expression of *CD9* and *CYP19A1*; ^31, 32^ and finally into syncytiotrophoblast (SCT) progenitors (characterized by expression of *GCM1* and of *TFAP2A/C*)^33, 34^ that will later fuse with the maternal decidua and become multinucleated. The multinucleated syncytium would not be captured due to their invasion of the maternal layer during dissection (Methods).

**Figure 2:**
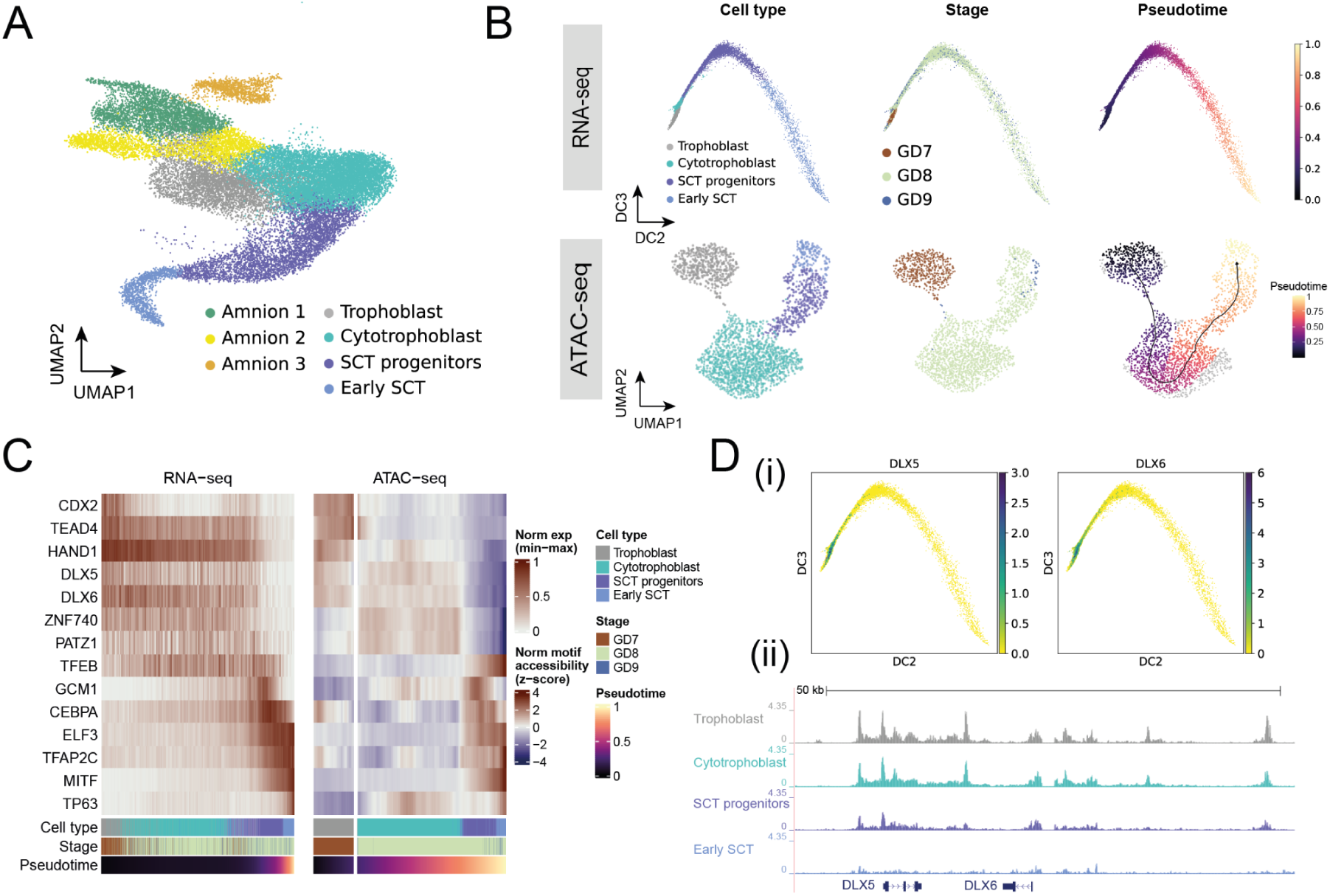
Chromatin accessibility and gene expression along the trajectory of early syncytiotrophoblast differentiation. **A)** Refined cell type annotation of extra-embryonic ectoderm cells. SCT; syncytiotrophoblast. **B)** Trophoblast cells/nuclei of the RNA-seq (top) and ATAC-seq (bottom) datasets are represented in low-dimensional embeddings which highlight a trajectory towards early syncytiotrophoblast. Cells are coloured by cell type (left), developmental time point (middle) and pseudotime (right), calculated independently for the RNA-seq and ATAC-seq atlases using diffusion pseudotime and ArchR trajectory inference respectively. The RNA-seq data is plotted with respect to the second and third diffusion components, whereas the ATAC-seq data is represented in a UMAP embedding. **C)** Heatmap of smoothed TF expression (left) and motif enrichment (right) along cells/nuclei of the RNA-seq and ATAC-seq SCT trajectories, ordered by their respective pseudotimes. The cell type, stage, and pseudotime value for each cell is indicated below. **D)** Expression of DLX5/DLX6 is restricted to trophoblast and cytotrophoblast cells, shown in the diffusion map embedding. (bottom) Genome browser view of the region surrounding the DLX5/6 locus, showing downregulation of accessibility along the differentiation trajectory.

To complement the transcriptional profiling of different trophoblast cell-types, and especially given the relatively limited information about trophoblast development in vivo, we next sought to elucidate the gene-regulatory programmes underlying trophoblast differentiation in the rabbit. To this end, we performed single-cell assay for transposase-accessible chromatin-sequencing (scATAC-seq) across the same time points for which scRNA-sequencing data was generated, resulting in chromatin accessibility profiles of 34,082 cells after quality control (Figure S5; Methods). Using the archR pipeline ^35^, cells were clustered and gene accessibility scores were used to perform cell type label transfer from the transcriptional atlas. Clusters containing cells of the different extraembryonic-ectoderm cell types were re-analysed and the cell type annotation was manually curated by inspection of chromatin accessibility of key marker genes and enrichment of transcription factors (TFs) motifs.

Ordering both the scRNA-seq and scATAC-seq data of the trophoblast-syncytiotrophoblast trajectory along pseudotime (Figure 2B) allowed us to compare the differential TF motif accessibility of known and putative regulators of trophoblast development and correlate this with differential expression of the matched scRNA-seq data (Figure 2C). Consistent with trophoblast development in human and mouse, we see enrichment for the binding motifs of TEAD4 ^36^ and CDX2 ^37^ at regions accessible during early trophoblast timepoints, which rapidly close as trophoblast cells mature. In cells assigned a cytotrophoblast identity, corresponding to the middle of the pseudotemporal ordering, we observed accessibility in regions associated with DLX5/6, ZNF740, and TP63 motifs. Later, during syncytiotrophoblast differentiation, we observed open chromatin in regions associated with the GCM1 motif ^38^, a known regulator of CEBPA and syncytin genes, the latter of which are directly involved in cell-cell fusion with the maternal decidua ^39, 40^. Finally, in the most mature syncytiotrophoblast cells, motifs associated with the binding of TFEB and MITF are highly accessible. These TFs are known to form homodimers or heterodimers, and are involved in vascularization and placental labyrinthine development ^41^. Overall, these patterns seen in rabbit in vivo are consistent with the limited information available for human, not only suggesting conservation of key temporal ordering of changes in chromatin accessibility during trophoblast development, but also establishing the first comprehensive in vivo dataset capturing this key process at single cell resolution for both transcription and open chromatin.

Interestingly, some patterns of changes in chromatin accessibility seen in the rabbit do not conform with previous mouse data; for example, we observed changes in chromatin accessibility at motifs associated with DLX5/DLX6 binding during cytotrophoblast differentiation, as well as corresponding expression (Figure 2D). These genes are known to be expressed in humans but are not expressed in murine trophoblast^42^. DLX5 and DLX6 are also known preeclampsia markers, with 69% of preeclamptic placentas showing upregulation in one study^42^. Moreover, these genes are implicated in proliferation of trophoblast and are normally downregulated during the differentiation process to syncytiotrophoblast. This conservation of markers between rabbit and humans reveals the rabbit may be a better model system for comparison to humans in preeclampsia studies

### A neighborhood-based comparison across species identifies rabbit-mouse differences in embryonic and extra-embryonic cell states and developmental trajectories

In the previous section, we focused on characterizing extraembryonic cell types that are present in the rabbit but which are underrepresented in mouse datasets at this stage of development. Next, we focused on cell types and developmental trajectories that are present across both species and examined which developmental processes were conserved and which showed divergence. To do this, we developed a computational approach to evaluate transcriptional similarity between k-nearest neighbor (kNN) graphs, constructed from scRNA-seq data of different species (Figure 3A). Independently for each species, we defined neighborhoods of cells across the entire kNN graph^43^ allowing us to locally approximate the transcriptional profile in precise regions of the gene expression manifold. In contrast to cell types or discrete clusters, neighborhoods are typically more granular, are independent of cell type annotations, and more optimally encapsulate gene expression changes, especially across continuous trajectories. Having defined neighborhoods within each species, we compute an average expression profile for each neighborhood before computing a correlation matrix (using orthologous genes) to measure the similarity between all pairs of neighborhoods across species (Methods).

**Figure 3:**
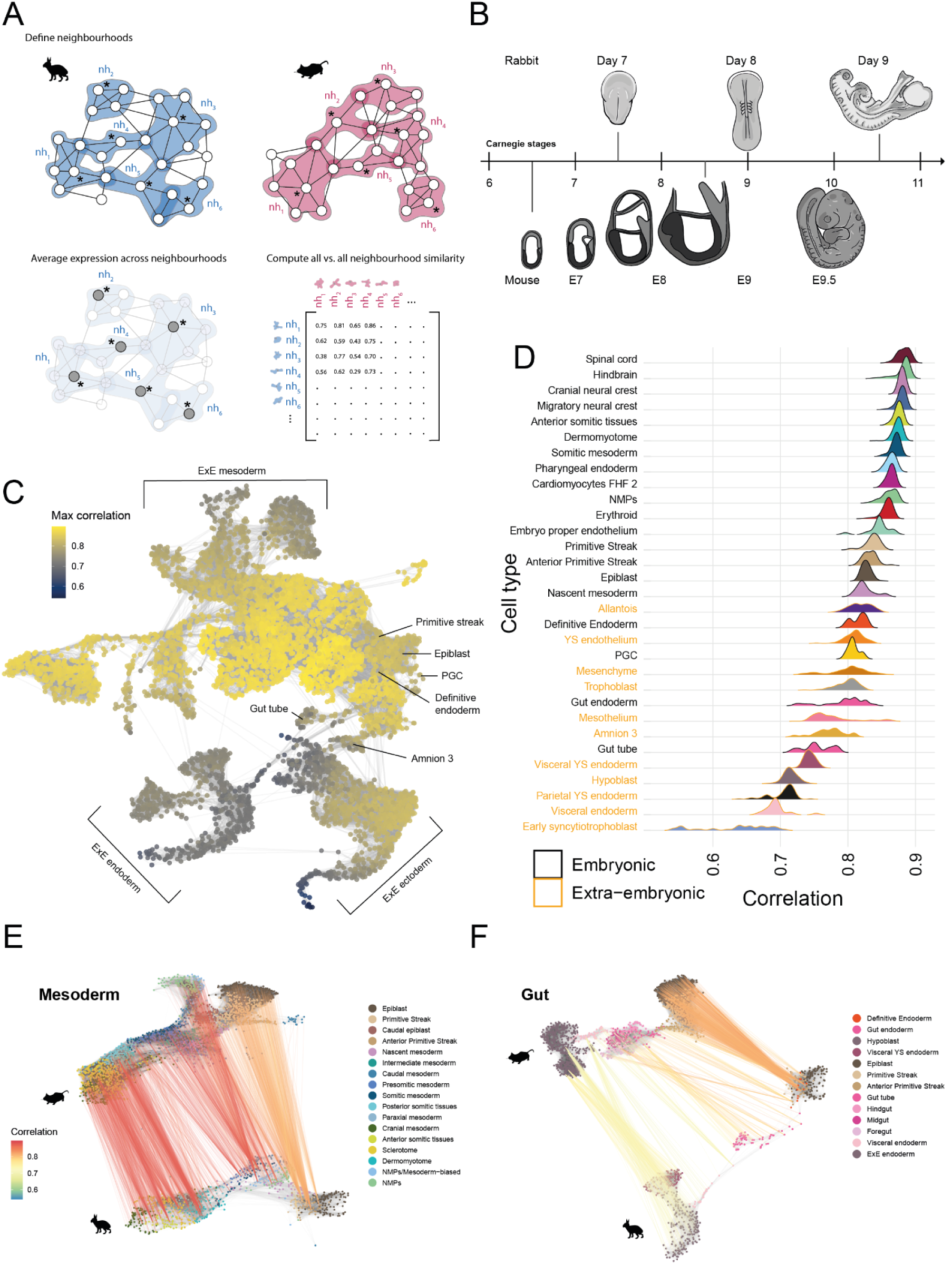
Rabbit and mouse neighbourhood comparisons identify regions of similarity and dissimilarity across species. **A)** Schematic of our neighbourhood comparison approach. Index cells (marked by *) are sampled across the kNN graphs of each species. The k-nearest neighbours of each index cell then collectively define overlapping neighbourhoods. The average expression profile within each neighbourhood is calculated and the correlation in expression across all neighbourhoods of both species is represented in a matrix of neighbourhood similarities. **B**) The timepoints sampled in both the rabbit and mouse ^26^ atlases are related according to Carnegie staging. **C)** Rabbit neighbourhoods, positioned with respect to the UMAP embedding of each index cell, are coloured by the maximum correlation value across all mouse neighbourhoods, highlighting regions of higher and lower similarity with the mouse. The underlying kNN graph is shown in grey. **D)** Maximum correlation scores are aggregated by the cell type of each neighbourhood index cell. The distribution of maximum correlation scores are shown, for a selected subset of cell types, in a ranked ridgeline plot. Extra-embryonic cell types are highlighted in orange. See a complete list in Figure S6B. **E, F)** A subset of rabbit and mouse neighbourhoods associated with mesoderm (**E**) and gut (**F**) differentiation trajectories are shown. Maximally correlated rabbit and mouse neighbourhood pairs (computed in both directions) are connected, where the line colour represents the value of maximum correlation.

We applied this approach to compare our transcriptional atlas of rabbit development with the E6.5 -E9.5 single-cell atlas of mouse development used in our cell type annotation ^26^. These timepoints overlap with our GD7, GD8 and GD9 rabbit samples, as assessed by Carnegie staging (Figure 3B). Using the matrix of correlations, computed across 5,253 rabbit and 14,034 mouse neighborhoods (Figure S6A), we first investigated which regions of the rabbit developmental landscape were conserved or divergent with the mouse. To do this, for each rabbit neighborhood, we extracted its maximum correlation value across all mouse neighborhoods (Figure 3C).

The most similar neighborhoods between species correspond to cell types within the embryo proper, particularly within the anterior and mid sections of the embryo (Figure S6D), including the neural crest, nervous system and mesodermal cell types (Figure 3D, S6B). By contrast, we noted that neighborhoods representing extraembryonic cell types, such as the amnion, parietal and visceral yolk sac endoderm, were among the least correlated cell types with the mouse, possibly reflective of the known morphological differences and divergent modes of development described previously, such as the depletion of Rauber’s Layer and generation of amnion from polar trophoblast. The Yolk Sac (YS) endothelium also shows a much lower maximum correlation distribution than the embryo proper endothelium (Figure 3D), potentially highlighting the impact of exposure to the extraembryonic environment.

Additionally, several embryonic cell types also showed relatively high divergence including the gut, primordial germ cells (PGCs) and cell types of the early GD7 embryo (Figure 3C,D; S6B,D). Interestingly, in the mouse, extraembryonic cell types play critical roles in the development of both the gut and the PGCs, via cell intercalation and signaling, respectively^44, 45^. In the former context, previous studies have shown that intercalated endoderm cells retain a transcriptional signature of their embryonic and extraembryonic origin^46^, indicating that differences in extraembryonic tissues may persist in cells of embryonic tissues. To investigate these observations in more detail, we examined how similarities between neighbourhoods vary along trajectories of differentiation. Specifically, we visualized mappings between maximally correlated neighborhoods along the reduced dimensional spaces of rabbit and mouse datasets for a subset of cell types relating to the development of specific lineages (Figure 3E-F, S7B-D). While neighbourhoods map very strongly across the whole spectrum of mesodermal cell types (Figure 3E), the trajectory of endoderm development is much less correlated between species (Figure 3F). As in Figure 3D, the extraembryonic component of the gut trajectory exhibits the lowest similarity between the rabbit and mouse, whereas neighbourhoods representing the epiblast and definitive endoderm, are more strongly correlated. Close to the point of intercalation, across neighbourhoods of the developing gut, the correlation is intermediary between values at the embryonic and extraembryonic origins, potentially as a result of cell mixing and convergence of transcriptional signatures. Moreover, many neighborhoods of the rabbit gut tube at GD9, have highest correlation with mouse neighborhoods at E8, suggesting a change in the timing of development (Figure S7A).

Taken together, our neighborhood-based analysis provides a general approach to compare single-cell RNA-seq datasets at a high level of granularity and independently of cell type annotations. Using correlations in expression, we are able to quantitatively assess differences between transcriptional profiles, which are often obfuscated, or difficult to interpret using integration-based methods. Applying this to the rabbit and mouse atlases we observe differences in extraembryonic tissues and in cell types such as the gut and PGCs, whose development is known to be influenced by extraembryonic structures and are linked to the maternal environment and cup vs disc embryo morphology ^47, 48^. We are also able to map neighbourhoods along continuous and discontinuous paths of differentiation, where we find varying dynamics in the conservation of cell states along different trajectories (Figure 3E-F, S7B-D). These results have implications for how we interpret observations from specific cell types and lineages in any one species and sets the scene for more extensive cross-species analysis when suitable datasets are reported for additional species.

### Leveraging the rabbit atlas to expand our understanding of early primate development

The quantity of cells, tissues and developmental stages captured in the rabbit and mouse atlases provide a comprehensive view into the development of these organisms. Given the scarcity of transcriptomic data from primate embryos, we next investigated whether the comprehensive mouse and rabbit resources could be leveraged, using automated cell type annotation tools, to gain deeper insight into the cellular makeup of sparse human and macaque datasets. Manually annotating cell types de-novo can be a challenging and time-consuming process, particularly when cell types are represented by small numbers of cells. In these cases, there is often little statistical power to confidently detect signals in marker gene expression above transcriptional noise. Being able to overcome these difficulties using model organism reference atlases will become increasingly relevant as more studies of early human development take place. To take advantage of our newly generated rabbit data, as well as the existing mouse datasets, we used SingleR to train cell type annotation models on the rabbit and mouse atlases. We then used these models to predict cell type annotations from the transcriptomic profiles of a representative human and macaque single-cell query dataset.

Tyser et al. 2021 reported a SMART-seq v2 dataset of 1,195 cells, obtained from a single CS7 human embryo between 16 and 19 days post-fertilisation ^25^. At this stage of development, the human embryo appears as a flat-disc, similarly to the rabbit embryo at day 7. Based on the transcriptomic profiles of these cells, we observe that our rabbit annotation model assigns several new cell type annotations, in addition to those consistent with the original publication (Figure 4A, S8A). For instance, cells originally classified as ‘hemogenic endothelial progenitors’ are subclassified into yolk sac endothelium, erythroid myeloid progenitors (EMP) and megakaryocytes (Figure 4A), which we were able to validate using known marker genes (Figure 4B). The model also identifies amniotic ectoderm cells and two PGCs. In the original study, these cell types were identified via a refined subclustering, requiring both complex computational analysis and a high-degree of domain expertise. Interestingly, the mouse-trained annotation model failed to classify these PGCs (Figure S8B). Given the relatively low neighborhood similarity scores between rabbit and mouse PGC neighborhoods, this suggests that the transcriptomic profile of rabbit PGCs are closer to that of the human. This is consistent with a recent study of PGCs in the rabbit which suggests that key regulators of PGC specification are shared across flat-disc species ^49^. Other differences are found in the prediction of epiblast cells, which the mouse-trained model labeled as ectoderm.

**Figure 4:**
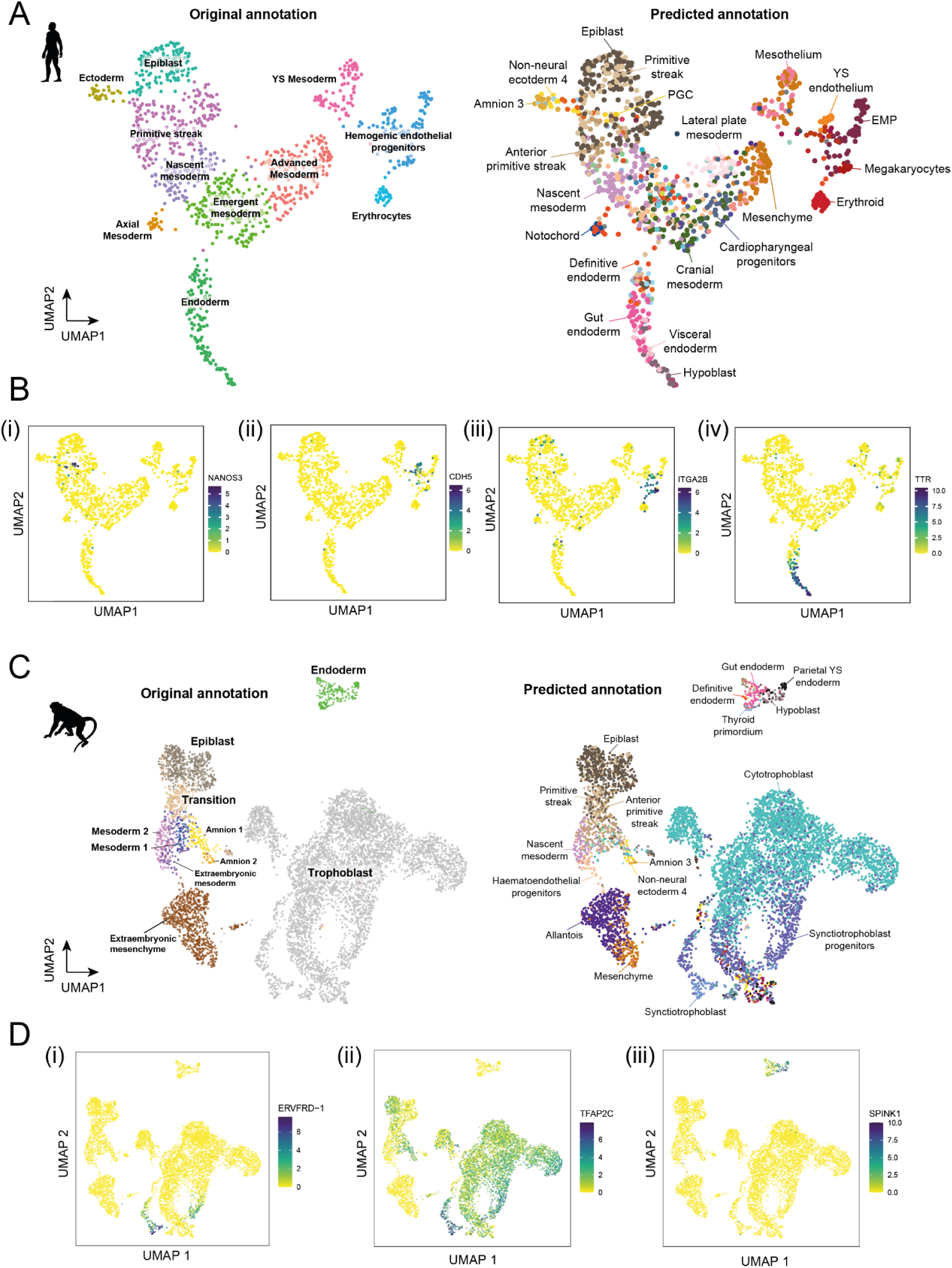
Automated annotation models trained on the rabbit atlas accurately classify cell types in sparse human and macaque data. **A)** UMAP of the Tyser et al. 2021 human embryo SMART-seq v2 data coloured according to the original cell type annotations (left) and the cell type labels predicted from training a SingleR model on the rabbit atlas (right). **B)** The rabbit model predictions are consistent with the expression patterns of known i) PGC, ii) endothelium iii) megakaryocyte and iv) extra-embryonic endoderm marker genes.

We applied the same approach to a single-cell RNA-sequencing dataset of cultured in-vitro macaque embryos spanning embryonic days 10, 12 and 14 ^50^. These stages of development correspond with Carnegie stages 5-6 ^51^ reflecting an earlier developmental stage than captured in our rabbit dataset, although partially overlapping with the E6.5-E7.5 stages of the mouse. Despite this, we find that both mouse and rabbit annotation models are able to replicate and refine the major cell types annotated in the original publication (Figure 4C, S8C,D). In addition to the epiblast, primitive streak, nascent mesoderm and amnion cells, which are concordant with the original labels, both rabbit and mouse trained models separate cells of the ‘extraembryonic mesenchyme’ cluster into allantois and mesenchyme annotations and distinguish gut endoderm cells from parietal and extraembryonic endoderm. Furthermore, the rabbit model predicts two domains of syncytiotrophoblast and syncytiotrophoblast progenitors within the original trophoblast labeled cluster. These regions overlap with the expression of the macaque-unique syncytin gene, *ERVFRD-1* and *TFAP2C* (Figure 4Di,ii). Given that implantation occurs around day 9.5-10.5 in the macaque ^52^, immediately prior to the timepoints represented here, it is possible that the trophoblast is transitioning through a similar differentiation process as observed in our rabbit atlas at GD8.

The ability of the mouse and rabbit references to precisely and accurately annotate distinct cell types within the human and macaque datasets illustrates the utility of using a more comprehensive reference set when performing cell type annotation. Since the rabbit and mouse cell types were annotated consistently, both rabbit and mouse annotation models provide unique insights, which can either increase confidence in the annotation or highlight possible cross-species differences. While the rabbit-trained model more accurately identified human PGCs and epiblast cells, the mouse-trained model predicted a smoother transition of mesodermal cell types across the UMAP embedding, possibly reflective of the higher resolution of 6-hour timepoints sampled (Figure S8B). This emphasizes the strengths of using both atlases, which we have made available (see Data Availability) to facilitate the annotation of cell types in other studies of mammalian embryogenesis.

### Rabbit Primitive Blood Niche Informs Human Haematopoietic Culturing Conditions

In the previous sections we have shown that the rabbit is an especially advantageous model system for studying extra-embryonic tissues, facilitating ready access to cell types that are not easily studied in the mouse. Amongst these cell types, our transcriptomic atlas contained a large number of yolk sac mesothelium cells, which have not been studied in great detail. This is of particular interest since the rabbit yolk sac is an important site for haematopoiesis and early fetal nutrition when the chorioallantoic stalk is not yet formed. The yolk sac consists of yolk sac endoderm, primitive blood cells, endothelium, and surrounding mesothelium. The rabbit yolk sac is particularly accessible compared to other mammals due to superficial implantation. Its large size also makes it easy to work with and characterize. Given the key role the yolk sac plays in early development, we elected to both characterize this developmental process and to interrogate how different cell types may interact with one another.

To this end, we mined our sc-RNAseq dataset for markers genes that are key to distinguish the different layers of the yolk sac hematopoietic niche in rabbit at GD9, and spatially resolved their localisation in situ, using single-molecule resolution microscopy (RNAscope) hybridisation. Using highly-spatially resolved RNAscope experiments, we identified blood cells, which can be seen inside of vessels formed by CDH5+ endothelium (Figure 5A). Some of the blood cells are RUNX1^-^ while others are RUNX1^+^. AHNAK^high^ mesothelium cells can be seen surrounding these CDH5+ endothelium cells (Figure 5A), with the blood cells in the center. The layer of large AHNAK^low^ endodermal cells matching the morphology of mouse endoderm cells can be seen opposite of the AHNAK^high^ layer. Visual inspection of these layers revealed that this basic structure commonly occurs in pairs, associated with each other via the endoderm-containing side, thus forming a mirrored bilayer structure constituting the rabbit blood islands (Figure 5B).

**Figure 5:**
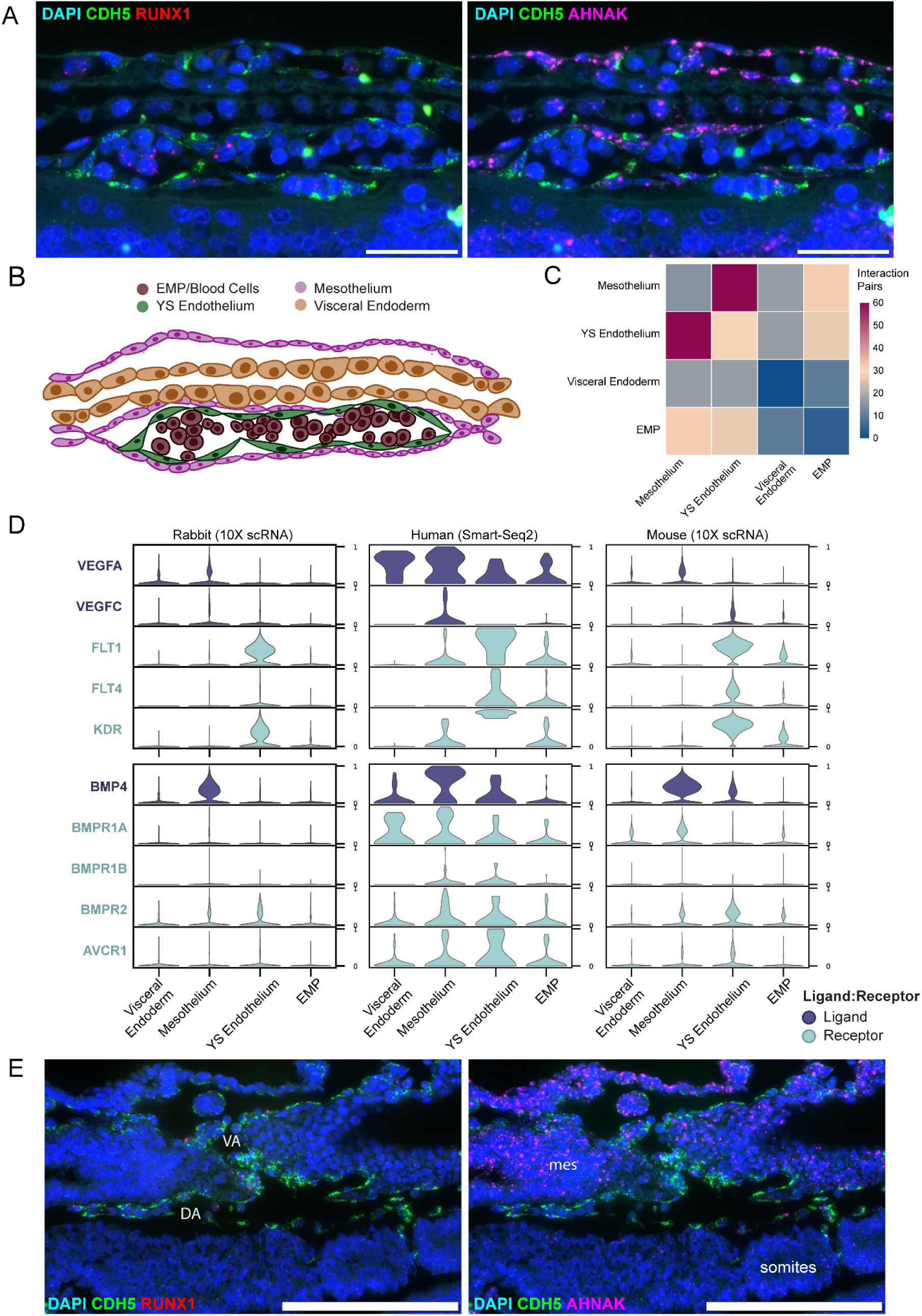
Rabbit yolk sack haematopoiesis exhibits conserved molecular markers with human in vitro development models and reveals signaling role of mesothelium. **A)** RNAscope image of yolk sac hematopoiesis with DAPI nuclear staining with probes for CDH5 and RUNX1 expression (left) and CDH5 and AHNAK expression (right). Cells are distinguished with RUNX1^+^ blood, CDH5^+^ endothelium, AHNAK^+^ yolk sac mesothelium and AHNAK^-^/CDH5^-^ visceral yolk sac endoderm. **B)** Schematic of the rabbit yolk sac haematopoietic niche according to the RNAscope images in A). **C)** Heatmap of CellPhoneDB interaction pair counts across different cells located within the yolk sac blood islands **D)** Violin plots of ligand-receptor expression across the yolk sac blood island cells for signaling involved in haematopoiesis such as VEGF/VEGFR (FLT1, KDR), and BMP/BMPR across the rabbit transcriptional atlas (left), CS7 human gastrula (Tyser et al. 2021, middle), and extended mouse transcriptional atlas (Rosshandler et al., manuscript in preparation, right). The ligands are colored in dark blue-purple, while the receptors are colored in cyan. **E)** RNAscope images for CDH5, DAPI, RUNX1, and AHNAK of the vitelline artery (VA) and dorsal aorta (DA) region with RUNX1+ CDH5+ presumptive haemogenic endothelium in the ventral portion of the dorsal aorta. mes = splanchnic mesenchyme.

The mesothelium has been overlooked as a potential signaling center for the maturation of blood cells in the yolk sac with the yolk sac endoderm thought to provide inductive signaling to the YS endothelium ^53, 54^. By leveraging the resolution of RNAscope, we observed that AHNAK+ cells are in direct contact with the YS endothelium. We also used CellPhoneDB ^55^ to predict the ligand-receptor interactions for these adjacent cell types. By inputting the cell-types that make up the yolk sac blood islands, we observed that the highest score of inferred interaction pairs are between mesothelium and YS endothelium (Figure 5E). This includes various extracellular matrix proteins like nectin, fibronectin-1 (FN1), collagen genes (Figure S9A,B) and important blood maturation ligands like VEGFA and VEGFC, expressed by the mesothelium. Having identified putative signaling crosstalk between rabbit mesothelium and endothelium, we next explored to what extent these interactions might be conserved in human, taking advantage of our refined annotation of the human CS7 dataset (Figure 4A). This analysis revealed high expression of VEGFA/C signaling from human mesothelium and consequent VEGFR1(FLT1) and FLT4 expression in the YS endothelium, thus supporting the interaction discovered in our rabbit data (Figure 5D)^56^. Of note, addition of VEGF was previously shown to promote in vitro primitive blood maturation from human pluripotent cells, yet the in vivo source of the cytokine had remained a mystery^57^. Looking into other regulators, we see that BMP4 is abundantly expressed in the mesothelium relative to the other cell types in the haematopoietic niche (Figure 5D). Given previous reports on BMP4 mediated induction of haematopoietic development ^57^, these results highlight the key role mesothelium may play in the yolk sac hematopoietic niche and similarities between rabbit and in vitro human blood culturing conditions.

In contrast to the mesothelium, analysis of genes that characterize the visceral yolk sac endoderm revealed hits related to cholesterol efflux and transport, indicating the endoderm might primarily facilitate provision of nutrition and metabolism to the embryo rather than being directly involved in blood maturation signaling^58^ (Figure S9B). Of note, high resolution RNAscope also reveals clusters of RUNX1+ CDH5+ haemogenic endothelial cells in the ventral side of the emerging dorsal aorta suggesting that an early wave of hemogenic endothelium may foreshadow the subsequent development of haematopoietic stem cells in this anatomical location (Figure 5E).

## DISCUSSION

In this study, we report high-resolution morphological and molecular maps of early rabbit development, covering gestational days 7, 8 and 9. Using cutting-edge single-cell genomics approaches, we characterized the transcriptional and chromatin accessibility profiles of over 180,000 individual cells, isolated from whole rabbit embryos, across gastrulation and early organogenesis. To our knowledge, this represents the most comprehensive view into the molecular landscape of early non-rodent mammalian development to date.

We leveraged these datasets, applying a novel neighborhood-based approach, to identify what developmental processes were most conserved, at the molecular level, between the rabbit and the mouse. We observed substantial differences in molecular profiles between the gut, PGCs and extra-embryonic cell types where transcriptional differences may be associated with the different embryo topologies, maternal environments, and implantation strategies of the rabbit and mouse. Related to this, we identified specific differences in genes regulating implantation and trophoblast differentiation between the two species, including the use of the transcription factors DLX5/DLX6 in the rabbit, which is similarly detected in the human trophoblast and is associated with the pathogenesis of pre-eclampsia ^42^. These results suggest that the rabbit may be a more suitable model for studying the development of the trophoblast than the mouse, an observation that is especially important given the lack of good in vitro models.

In general, we find that the large size and late, superficial implantation of the rabbit embryo allows for the efficient capture of extra-embryonic tissues. Combining our transcriptomic atlas with RNAscope in-situ hybridisation, we are able to see that the yolk sac mesothelium is in direct contact with haemogenic endothelium and expresses respective ligand-receptor pairs of key blood maturation genes, such as VEGFA and VEGFC. These findings suggest that the mesothelium plays a more important role in hematopoietic signaling than previously recognised.

Altogether, our results highlight the utility of the rabbit as a model for a new wave of mammalian development research that will combine the power of traditional comparative embryology with state-of-the-art comparative single cell genomics approaches. When researching rabbit development for this manuscript, one of the most useful references turned out to be the 1905 ‘Plate’ of rabbit development, published as one of 16 “Normal Plates of the Development of the Vertebrates” edited by the German anatomist Franz Keibel (16 volumes, 1897–1938)^59^. It may have taken over 100 years, but it is now realistic to imagine a similar compilation of single cell genomics atlases for over a dozen vertebrates within the next few years. Coupled with innovation in experimental techniques such as embryo culture, genome modification and lineage tracking, single cell comparative genomics will then likely emerge as a new energizing force accelerating the use of model organisms to decipher early human development and drive advances in translational medicine.

## Methods

### Rabbit Embryo processing for scRNA-Seq

New Zealand White Rabbits (Oryctolagus cuniculus) from Envigo (RMS) UK Ltd, were mated at Labcorp Early Development Laboratories Limited (Eye, Suffolk; formerly known as Covance Laboratories Limited) by natural mating. After mating each female was injected intravenously with 25 i.u. luteinizing hormone. GD0 was defined as the day of mating. On GD7, 8 or 9 the pregnant rabbits were sacrificed, and the uteri were harvested and shipped fresh in Phosphate Buffered Saline (PBS) on ice to the Jeffrey Cheah Biomedical Centre. Embryos were dissected from the uteri at gestational day (GD) 7, 8, and 9 using a Leica brightfield microscope and fine point tweezers using 10% heat-inactivated Fetal Bovine Serum (FBS) in PBS. Embryos were selected based on morphology matching developmental day of dissection as described previously ^59^. Selected embryos or dissected structures (see Table S1) were rinsed in PBS, centrifuged for 100 x g for 3 minutes before being individually dissociated with TrypLE™ Express (Gibco™) by incubating for 6-10 minutes at 37°C with occasional agitation of the tube by flicking the base of the tube in order to ensure even dissociation of the embryo. Dissociation was quenched with 5mL 10% heat-inactivated FBS in PBS and filtered using a 30µm Sysmex CellTrics® filter. Cells were centrifuged for 300 x g for 3 minutes and resuspended in 0.04% BSA in PBS. Cells were filtered through a 40μm Flowmi tip strainer (VWR) to minimize volume loss during filtration and then counted using a haemocytometer.

Six GD7 embryos were dissociated separately, and processed as individual samples. Two pools of 3 GD8 embryos with no visible somites were processed, as well as 3 GD8 individually processed embryos with 4 somites apiece. Two GD9 embryos were split between embryo proper and extraembryonic tissues. Embryo proper portions were dissociated and split across two separate samples. Another 2 GD9 embryos were individually processed and split into the anterior, mid, posterior, and yolk-sac region and to provide spatial information. Pool sizes were chosen to maximize the number of cells recovered per sample while minimising the doublet rate. Embryos were partitioned if necessary across multiple samples to prevent exceeding the recommended amount of loaded cells.

Cell solutions and scRNA-seq libraries were processed by the CRUKCI genomics core facility using Single Cell Gene Expression v3 from 10X Genomics following manufacturer’s instructions. Samples were sequenced following manufacturer’s recommendations on an Illumina NovaSeq 6000 platform.

### Rabbit embryo processing for scATAC

Embryos dissected from the same time-points for transcriptional profiling were flash frozen by placing whole embryos into Corning® Cryogenic Vials and immediately submerging the vials in liquid nitrogen. Embryos were stored at -80C for later use. Nuclei were extracted following a 10X Genomics demonstrated protocol for nuclei isolation for single cell ATAC on frozen tissues (CG000212). For each timepoint, two GD7 embryos were pooled together for nuclear extraction, 1 GD8 embryo was split across two samples, and 1 GD9 embryo was split into the extraembryonic portion for one sample and the embryo portion was split across four samples. Frozen embryos were placed on ice with 500uL chilled 0.1X lysis buffer and homogenized using RNase-Free Disposable Pellet Pestles (Fisher Scientific). Samples were filtered using a 70 µm Flowmi Cell Strainer followed by a 40 µm Flowmi Cell Strainer. Nuclei were counted in a haemocytometer using Trypan Blue Solution 0.4% (Sigma-Aldrich, Cat. No. T8154-20ML) and were resuspended in 1X Nuclei Buffer (10X Genomics). Nuclei solutions and libraries were processed by the CRUKCI genomics core facility using Single Cell ATAC (v1.1) from 10X Genomics following manufacturer’s instructions. Samples were sequenced following manufacturer’s recommendations on an Illumina NovaSeq 6000 platform.

### Improving transcriptome mapping

After processing the single-cell RNA-sequencing data with Cell Ranger using the Ensembl OryCun2.0 rabbit reference transcriptome, we observed a low percentage of reads mapping to the transcriptome. Visualizing read coverage across the genome, we discovered that a large majority of the 10X sequencing reads were aligning to regions upstream of the 3’ end of annotated genes (Figure S2A-B). Furthermore, it was evident that several genes, which are known to be well-conserved, were missing from the rabbit annotation. To improve the transcriptome mapping, we extended the 3’ annotation of genes by 600bp and added human genes that aligned to unannotated regions in the rabbit genome.

The decision to extend genes by 600bp was determined by analyzing distances of intergenic reads from their nearest annotated gene (Figure S2A). Intergenic reads were extracted from the Cell Ranger BAM output file of the SIGAC11 GD8 (Table S1) sample by filtering on the ‘RE’ BAM alignment tag using SAMtools. This revealed that the majority of reads fell within 600bp from the 3’ end of the nearest gene, suggesting that a 600bp extension would provide a reasonable compromise between capturing mising reads and avoiding overlaps with nearby annotations. We also prevented this explicitly, ensuring that extensions were only added to each gene providing they did not overlap with an existing annotation. The GTF file of the rabbit transcriptome was modified by adding CDS and exon entries to ensure they were counted by the Cell Ranger pipeline.

To obtain a list of features that may be absent from the rabbit annotation, we identified genes that have a one-to-one orthology relationship between the mouse and human but were missing from the rabbit reference. Given the human gene ID for each feature, we located positions in the human genome relating to transcripts of that gene and specifically, the positions of each transcript’s 3’ most exon and UTR. Using the Ensembl Compara Perl API, we then obtained alignments for these sequences in the rabbit transcriptome, taking only those with maximum alignment scores. Exon and CDS annotations spanning these positions were then added to the rabbit GTF file, similar to the extended 3’ ends. Additional measures were taken to ensure that the alignments extracted did not overlap existing annotations (including the added extensions) and that alignment sequences for the same transcript were proximal (i.e. not mapping to different chromosomes).

As a result of these changes, the median number of genes and UMIs increased from 1875 to 2661 and from 6987 to 10,126 respectively. Moreover, 1648 additional genes were added from the alignments, which included several known marker genes, including FOXC1 and SIX3 (Figure S2D). Together these improvements made a noticeable impact on the quality of data used downstream (Figure S2C,E).

### scRNA-seq preprocessing via Cell Ranger

The FASTQ sequencing files were processed with Cell Ranger 3.1.0 using default mapping arguments. Reads were mapped to the OryCun2.0 genome from Ensembl with the modified 3’ extension GTF annotation file mentioned previously.

### Swapped molecule removal

Swapped molecular counts were corrected as previously described (Pijuan-Sala et al. 2019). Briefly, molecule counts that were derived from barcode swapping were removed from all samples by applying the DropletUtils function ‘swappedDrops’ (default parameters) to groups of samples that were multiplexed for sequencing.

### Cell calling for scRNA-Seq

Cells were called as adapted from (Pijuan-Sala et al. 2019). Briefly, cell barcodes that were associated with real cell transcriptomes were identified using emptyDrops^60^, which assesses cells with RNA content distinct from ambient background RNA, the latter determined from barcodes associated with fewer than 100 unique molecular identifiers (UMIs). Cells with P < 0.01 (Benjamini–Hochberg-corrected) and at least 3500 UMIs and 900 unique genes were considered for further analysis.

Additionally, cells with mitochondrial gene-expression fractions greater than 24.03% were excluded. Rabbit mitochondrial fractions were suspected to be higher than mouse due to species differences or genome annotation. The thresholds were determined by considering a median-centred median absolute deviation (MAD)-variance normal distribution; cells with mitochondrial read fraction outside of the upper end of this distribution were excluded (where outside corresponds to *P* < 0.05; Benjamini–Hochberg-corrected).

### Doublet removal for scRNA-Seq

Doublets were scored as previously described ^10^. First, a doublet score was computed for each cell by applying the ‘doubletCells’ function (scran R package) to each sample separately. This function returns the density of simulated doublets around each cell, normalized by the density of observed cell libraries. High scores indicate high doublet probability. We next identified clusters of cells in each sample by computing the first 50 principal components (PCs) across all genes, building a shared nearest-neighbour graph (10 nearest neighbours; ‘buildSNNGraph’ function; scran R package), and applying the Louvain clustering algorithm (‘cluster_louvain’ function; igraph R package; default parameters) to it. Only HVGs (calculated separately for each sample) were used for the clustering. This procedure was repeated in each identified cluster to break the data into smaller clusters, ensuring that small regions of high doublet density were not clustered with large numbers of singlets. For each cluster, the median doublet score was considered as a summary of the scores of its cells, as clusters with a high median score were likely to contain mostly doublets. Doublet calls were made in each sample by considering a null distribution for the scores using a median-centred MAD-variance normal distribution, separately for each sample. The MAD estimate was calculated only on values above the median to avoid the effects of zero-truncation, as doublet scores cannot be less than zero. All cells in clusters with a median score at the extreme upper end of this distribution (Benjamini–Hochberg-corrected *P* < 0.1) were labelled as doublets. A final clustering step was performed across all samples together to identify cells that shared transcriptional profiles with called doublets, but escaped identification in their own samples. Clusters were defined using the same procedure as was applied to each sample, with the exceptions that sub-clustering was not performed, and batch-corrected principal components were used (see ‘Batch correction’, above). To identify clusters that contained more doublets than expected, we considered for each cluster the fraction of cell libraries that were called as doublets in their own samples. We modeled a null distribution for this fraction using a median-centered, MAD-estimated variance normal distribution as described for the median doublet score in each sample, above, and called doublets from the distribution as in each sample, above.

### Stripped nucleus removal for scRNA-Seq

Cells with considerably lower mitochondrial gene expression and smaller total UMI counts compared with other clusters were removed as previously described (Pijuan-Sala et al. 2019). We assumed that these clusters consisted of nuclei that had been stripped of their cytoplasm in the droplets, and therefore excluded them from downstream analyses.

### Normalization for scRNA-Seq

Transcriptome size factors were calculated as previously described (Pijuan-Sala et al. 2019) using ‘computeSumFactors’ from the scran R package. Cells were pre-clustered with the ‘quickCluster’ function using the parameter ‘method=igraph’ (using the scran R package), and minimum and maximum cluster sizes of 100 and 3,000 cells, respectively. Raw counts for each cell were divided by their size factors, and the resulting normalized counts were used for further processing.

### Batch correction for scRNA-Seq

Batch effects were removed using the ‘fastMNN’ function from the batchelor R package. The top 3000 HVGs were calculated with the ‘modelGeneVar’ and ‘getTopHVGs’ functions in ‘scran’. These HVGs were then used by fastMNN to compute the top 50 principal components. To ensure that the correct mutually nearest neighbours were identified, we enforced a particular order in which to combine samples, using the ‘merge.order’ parameter of ‘fastMNN’. Specifically, we merged samples in reverse order of developmental stage and in decreasing order of the number of cells within each timepoint. For the GD9 samples, we took additional care to merge samples of the same anatomical dissection before those from different parts of the embryo. As a result of this step, all 26 samples were integrated into a combined principal component space. These principal components were used for all downstream analysis steps (e.g, constructing nearest-neighbour graphs).

### Visualization and clustering

A UMAP embedding of the whole dataset was computed using Scanpy (‘scanpy.api.tl.umap’)^61^. The 300 nearest neighbors in the batch-corrected principal component analysis were considered, with a ‘min.dist’ parameter of 0.9. Force-directed graph layouts were also computed on the batch corrected PCs with the ‘scanpy.api.tl.draw_graph’ function using the ForceAtlas2 algorithm.

The whole dataset was clustered using the Leiden algorithm (scanpy.api.tl.leiden) using the same neighborhood graph constructed for the UMAP embedding. Clusters were generated using a range of resolution parameters (1 - 10) to assist in making cell type annotations at varying levels of coarseness. We also performed clustering within known lineages and subsets of the data to capture different sources of variation.

### Cell type annotation

Cell type annotations were initially predicted at single-cell resolution by training an automated annotation model on a recent single-cell atlas of mouse gastrulation and early organogenesis ^26^. The Rosshandler dataset consists of 430,339 cells across embryonic days 6.5 to 9.5, sampled at 6-hour intervals. These cells were manually curated into 87 different cell types following consultations with domain experts and a detailed examination of known marker genes. Given that the rabbit is less-well studied and has fewer known marker genes than the mouse, transferring labels across species served to provide a fast, initial annotation that could be scrutinized and validated in more detail.

To transfer annotations from the mouse to the rabbit, we utilized SingleR, an automated annotation method that assigns cell type labels based on correlation in expression between reference and query cells ^62^. In order for the cell type annotation model to relate the rabbit and mouse datasets, we constructed a common feature set between the atlases using one-to-one ortholog genes. For each rabbit gene, we extracted its mouse homolog from Ensembl and cross-referenced it with the Ensembl codes of the mouse atlas. Many-to-one and many-to-many genes were excluded by filtering on the orthology type also provided by Ensembl.

We then trained a SingleR model on the mouse atlas (with ‘trainSingleR’ from the ‘SingleR’ package using parameters ‘de.n = 50’ and ‘de.method = Wilcox’), providing the cell type annotations from the original study. Given the large number of cells, we opted for the pseudobulk option within SingleR, which clusters cells of the same cell type into pseudo-samples, reducing the computational work.

The cell type predictions from SingleR assisted in assigning identities to each cluster. Since the annotation model makes predictions independently for each query cell, clusters with a dominant cell type prediction were suggestive of a reliable annotation. We could also assess the assignment confidence for each cell using the ‘delta’ values provided by SingleR, which quantify the difference between the assigned annotation and the mean across all other possible annotations. To validate the cell type predictions we also relied on timepoint information, spatial information from the GD9 samples and the expression of cell type marker genes. In some cases, such as in the annotation of parietal endoderm, we also visualized the expression of marker genes specific to a given cluster, using RNAscope. All of these different sources of information were considered in order to assign cell type labels to clusters.

### Rabbit-mouse integration

The rabbit and mouse ^26^ datasets were integrated into a joint embedding using SAMap version 0.1.6 ^63^. An initial mapping between rabbit and mouse features was first obtained by reciprocal BLAST aligning the rabbit and mouse transcriptomes. The transcriptomes were obtained through Ensembl and aligned using the ‘map_genes.sh’ script, provided with SAMap (https://github.com/atarashansky/SAMap/). This script outputs a table of sequence similarity scores between rabbit and mouse transcripts, which are used by SAMap as an initial weighting between features. Since the features of the rabbit and mouse datasets are given in terms of Ensembl gene IDs, prior to running SAMap, we linked each Ensembl transcript ID with its associated gene ID. The SAMap algorithm also utilises self-assembling manifolds ^64^ to align datasets and so these were computed for the rabbit and mouse atlases using Scanpy (scanpy.tl.external.sam). We then constructed a SAMAP object, providing the rabbit and mouse SAM objects, the transcript to gene mappings and the map_genes output directory. We then ran the SAMap pipeline on this SAMAP object (SAMAP.run) to integrate the two datasets. For visualisation purposes, we re-computed a UMAP embedding on the integrated anndata object (SAMAP.adata) using a minimum distance parameter of 0.8.

### RNA-seq trophoblast analysis

Cells annotated as trophoblast, cytotrophoblast, syncytiotrophoblast progenitors and early syncytiotrophoblast were isolated for trajectory analysis. A diffusion map low-dimensional embedding^65, 66^ was computed (using ‘scanpy.api..tl.diffmap’) with 30 components on the batch corrected kNN graph. The GD7 trophoblast cell with highest value in its second diffusion component was then chosen as the root cell for computing diffusion pseudotime^67^ (with scanpy.api.tl.dpt) on the 30 diffusion components, specifying 0 branchings.

In Figure 2C, cells were ordered according to diffusion pseudotime. Expression values for each gene were then smoothed along this pseudotemporal ordering by calculating a moving average with a window size of 49 cells. These smoothed values were normalised to between 0 and 1 using min-max scaling. The same procedure was used to plot the ATAC-seq motif accessibility values, except that cells were ordered along pseudotime calculated in ArchR (see ‘scATAC-seq trophoblast analysis’) and a z-score normalisation was applied in replace of min-max scaling, to account for positive and negative accessibility scores. The combined heatmap was plotted using the ‘ComplexHeatmap’ package in R.

### scATAC-seq pre-processing

FASTQ files of scATAC-sequencing were mapped to the rabbit genome (OryCun2.0) using cellranger-ATAC version 2.0.0. Full analysis is performed using the ArchR pipeline^35^, where reads on unplaced scaffolds in the rabbit genome were ignored in the pipeline. Arrow files were created from CellRangers fragment files using createArrowFiles, using minTSS=2 and minFrags=3000, followed by doublet detection and removal using addDoubletScores with k=15. After inspecting quality distributions, new thresholds were determined (minTSS=2.9 and minFrags=8000). A total of 34,082 cells were used in the analysis. Cell numbers and quality statistics per sample can be found in Table S1 and Figure S5, respectively.

### scATAC-seq dimensionality reduction and peak calling

The TileMatrix, containing read counts per 500 bp bins of the entire genome, was used for a first round of dimensionality reduction using addIterativeLSI, using dimsToUse=1:45 and nFeatures=60000 and clustering using addClusters with resolution=1. These clusters were then used to create ‘Pseudo-Bulk Replicates’ using addGroupCoverages with useLabels set to TRUE. These Pseudo-Bulk Replicates were subsequently used for peak calling with Macs2 ^68^ (version 2.2.7.1) using addReproduciblePeakSet, with genomeSize=2.7e9, cutOff=0.001, and extendSummits=250, resulting in a final set of 332,773 peaks. The detected peaks were then used to create the PeakMatrix, containing read counts per peaks. The second round of dimensionality reduction and clustering was performed on the PeakMatrix, using the same parameters as for the TileMatrix. UMAP was performed with addUMAP, using nNeighbors=45 and minDist=0.5.

### scATAC-seq and scRNA-seq integration

To perform cell-type annotation of the scATAC-seq cells, integration was performed with the labeled scRNA-seq dataset for each stage separately. For each stage, dimensionality reduction was performed using addIterativeLSI, using dimsToUse=1:35 and nFeatures=55000 and clustering was performed using addClusters with resolution=1.5 to create high-resolution per stage clusters. Imputation matrices were added using addImputeWeights. Next, scRNA-seq integration was performed using addGeneIntegrationMatrix, which compares the estimated gene activity stored in the ‘GeneScoreMatrix’ for scATAC-seq with the gene expression of the scRNA-seq dataset in order to perform label transfer for cell-types and add a ‘GeneIntegrationMatrix’ containing integrated RNA expression counts.

### scATAC-seq motif deviations

For each peak, TF motifs were detected using addMotifAnnotations, searching for motifs in the ‘cispb’ set for Homo Sapiens, as the motifs between human and rabbit are expected to be highly conserved. To compute motif deviations, first a set of background peaks were detected using addBgdPeaks, followed by generation of the ‘MotifMatrix’ by addDeviationsMatrix, which stores the deviation and Z-scores for each motif per cell.

### scATAC-seq trophoblast analysis

Trophoblast and amnion cells were isolated from the total dataset by isolating cells belonging to clusters 1 to 5. To manually curate the annotation of these cells, dimensionality reduction and reclustering was performed with addIterativeLSI, using dimsToUse=1:25 and nFeatures=25000 and addClusters with resolution=1. Markers of the ‘GeneScoreMatrix’ and ‘MotifMatrix’ per cluster were used to relabel Trophoblast, Cytotrophoblast, Syncytiotrophoblast progenitors, Syncytiotrophoblast, and Amnion cells. Next, Amnion cells were excluded and dimensionality reduction was repeated on these cells with dimsToUse=1:10 and nFeatures=15000, and the UMAP was generated using nNeighbors=35 and minDist=0.8. Trajectory analysis was performed by determining pseudotime ordering using addTrajectory over the UMAP embedding. Motifs with variable accessibility across pseudotime were subsetted to those of particular interest, which were correlated with TF expression in the scRNA-seq dataset across trophoblast celltypes, or which are known from the literature.

### Neighbourhood comparisons

To define neighbourhoods, we used the same construction employed for differential abundance testing ^43^. Independently for the rabbit and mouse, the top 50 PCA components were used to construct a kNN graph (with k=30 neighbors). Neighbourhoods were then defined by aggregating the k-nearest neighbours of a randomly sampled set of index cells. The final set of neighbourhoods were refined from this initial selection to prevent oversampling and to create larger, more representative neighbourhoods (see ^43^ for details). These steps were performed by the ‘buildGraph’ and ‘makeNhoods’ functions from the ‘miloR’ package using a sampling proportion of 0.05. From this we obtained 5,253 rabbit and 14,034 mouse neighborhoods with a mean neighbourhood size of 104.8 and 134.4 cells respectively (Figure S6A).

We next identified a set of features with which to compare neighbourhoods across the two species. Specifically, we selected the intersection of the top 2000 highly variable genes, computed independently for each dataset. The intersection of genes was chosen to avoid confounding differences in expression with technical variation resulting from the lower quality rabbit genome annotation. It ensured that genes selected for comparisons were expressed and highly variable across both datasets. We also experimented with the number of highly variable genes but found that our results changed very little above 2000 HVGs. Features were selected using the ‘getScranHVGs’ function from the scran R package. We excluded from this mitochondrial genes and those associated with cell cycle GO terms. Finally we selected only those which are one-to-one orthologs across the rabbit and mouse. Our final set of features consisted of 796 genes.

Using this set, we computed the mean expression profile within each neighbourhood (using the normalised and log-transformed gene counts). We also implemented a version of gene-specificity, used in previous cross-species comparisons, to account for differences in quantification and absolute values between datasets ^69, 70^. Specifically, the within-neighbourhood averages for each gene (*g_i_*) were scaled by the mean across all neighbourhoods (*i* ∈ {1, … *N*}).

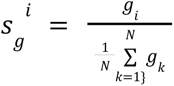

The gene specificity values 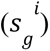 for each gene, across all neighbourhoods, were then used to compute the Pearson correlation between all pairs of rabbit and mouse neighborhoods forming a matrix of neighbourhood similarity.

In Figures 3 and S7, we visualised neighbourhood similarities across the rabbit and mouse UMAP embeddings. For each rabbit neighbourhood we extracted its maximum similarity score with any mouse neighbourhood (and vice versa). We then plotted each neighbourhood according to the UMAP position of its index cell, and coloured each point by the neighbourhood’s maximum correlation value. In Figures 3D, the maximum correlation values are aggregated according to the cell type annotation of each index cell, to obtain a distribution of similarity scores across each cell type. Finally, for the trajectory comparisons, subsets of neighbourhoods were extracted from both the rabbit and mouse datasets whose index cell was annotated as one of a specified set of cell types. These neighbourhoods were then plotted in the same way as Figure 3C and repositioned or reflected to facilitate a clear visual comparison. Lines were drawn between maximally correlated neighbourhood pairs, computed in both the rabbit-mouse and mouse-rabbit directions. The line colour indicates the strength of correlation for each mapping.

### Primate cell type annotation

In our cell type annotation analysis we utilised the Tyser et al. 2021^25^ and Yang et al. 2021 ^50^ single-cell datasets of early human and macaque development. The Tyser et al. 2021 SMART-seq dataset consists of 1,195 cells from a single CS7 human embryo acquired through the Human Developmental Biology Resource. We downloaded the normalised gene expression values and UMAP coordinates from http://human-gastrula.net/. Conversely, the Yang et al. 2020 dataset was generated through 10X genomics single-cell RNA-sequencing of in-vitro cultured macaque embryos, at days 10, 12 and 14. We accessed raw count data through the GEO database under accession number GSE148683 and downloaded associated metadata from the accompanying shiny app at https://www.nhp-embryo.net/. To combine the day 10, 12 and 14 samples, we performed batch correction with fastMNN with k=20. We also normalized and log-transformed the raw counts using ‘computeSumFactors’ and ‘logNormCounts’ functions in the ‘scran’ and ‘scuttle’ R package.

To predict cell types in the human and macaque datasets, we applied the same approach used to annotate the rabbit atlas originally. We trained SingleR annotation models on the rabbit and mouse datasets both jointly and independently. Datasets were filtered to include only one-to-one ortholog genes between each reference (rabbit or mouse) and query (human or macaque) dataset. As before, the training and classification steps were performed using the ‘trainSingleR’ and ‘classifySingleR’ functions from the ‘SingleR’ R package, with the ‘aggr.ref=TRUE’ option. In Figure 4, we display the ‘pruned’ cell type annotations from SingleR, which have undergone a filtering step to remove low-quality assignments.

### Histology

Timed mating: Male/female ratio 1:1. The female and male mated twice (morning and afternoon). Post-mating, the female rabbit was injected intravenously luteinising hormone (dose 25 i.u., Luveris (75 IE, Merck)). The day of mating was counted as GD 0. Embryo collection and processing: Pregnant rabbits from timed matings were euthanized at gestational days (GD) 7, 8 and 9 by an i.v. administration of a lethal dose of pentobarbital (0.35 ml/kg, Euthasol (400 mg, Dechra Veterinary Products A/S)). The uteri were dissected out and transferred to ice-cold phosphate buffered saline. Some embryos were dissected out while others were left intact in uterus to preserve extraembryonic structures. Dissected tissues were transferred to 10% neutral buffered formalin then allowed to fixate for 48 to 96 hours. After fixation the specimens were processed in an automated tissue processor (Leica ASP300S) and embedded in paraffin.

The paraffin embedded specimens were serially sectioned on an AS-410M fully automatic microtome (Axlab) producing 6µm sections spanning the entire embryo in each specimen. Hematoxilin and eosin staining: Slides were deparaffinized with xylene and rehydrated to water. Mayer’s hematoxilin solution (Sigma-Aldrich, MHS80) was applied for 5 minutes followed by washing in tap water for 5 minutes. Eosin (Sigma-Aldrich, HT110280) was then applied for 5 minutes followed by washing and dehydration in a graded ethanol series to xylene. The slides were mounted with pertex and whole slide scanned on an Olympus VS200 scanner using a 20x NA 0.8 objective.

### RNAscope

mRNAs were detected using the automated RNAscope LS Multiplex Fluorescent Assay on a Leica Bond RX autostainer with the following sets of probes: TFAP2C-C1, POU5F1-C2, NANOS3-C3 (ACD, Cat. No.: 1103588-C1, 513278-C2, 1103578-C3), CDH5-C1, RUNX1-C2, AHNAK-C3 (ACD, Cat. No.: 1043138-C1, 1043148-C2, 1043158-C3) and LGALS2-C1, DKK1-C2, SERPINC1-C3 (ACD, Cat. No.: 1092968-C1, 1092978-C2, 1092988-C3). The slides were first deparaffinized (Leica Biosystems, AR9222) followed by pre-treatment with BOND Epitope Retrieval Solution 2 (Leica Biosystems, AR9640) and protease (ACD, 322800) followed by probe hybridization. The probes were detected using the RNAscope LS Multiplex Reagent Kit (ACD, Cat. No. 322800) and Opal 520 (C1), Opal 570 (C2) and Opal 690 (C3) fluorophores (Akoya Biosciences, 1487001KT, FP1488001KT, FP1497001KT). The slides were counterstained with DAPI (Sigma-Aldrich, D9564), mounted with Prolong Diamond antifade mountant (Thermo Fisher Scientific, P36970) and whole slide imaged on an Olympus VS200 slide scanner equipped with DAPI, FITC, Cy3 and Cy5 filter sets using 20x NA 0.8 or 40x NA 0.95 objectives. Images were then processed for publication using olympusVIA.

### CellphoneDB analysis of Yolk Sac Haematopoiesis

Cell-cell interactions were predicted using a previously curated list of ligand-receptor and receptor-receptor pairs using CellphoneDB v.2.0 ^55^. Due to the exponential nature of possible interactions across the full dataset, only a subset of cell types corresponding to the rabbit yolk sac and blood islands were used, covering ‘EMP’, ‘visceral endoderm’, ‘yolk sac endothelium’, and ‘YS mesothelium’. The visceral endoderm cell type in the CellphoneDB heat map analysis used the merged cell type of ‘visceral endoderm’, ‘visceral ys endoderm 1’, and ‘visceral ys endoderm 2’ to cover all possible interaction pairs between different subtypes of the visceral endoderm as well as other cell types.

### Data and code availability

Raw sequencing data are available on ArrayExpress with the following accessions: scRNA-seq: E-MTAB-11836; scATAC-seq: E-MTAB-11804. Links to the data and code are available at https://marionilab.github.io/RabbitGastrulation2022/. The web page also links to processed single-cell transcriptomics data in a variety of formats for loading into both R and python analysis pipelines. The data is also available to explore interactively via a web app accessible through the same link. All other data are available from the corresponding authors on reasonable request.

## Acknowledgements

We would like to thank K. Katarzyna and the CRUK-CI Genomics Core for their help with the 10X Genomics sample processing; R. Argelaguet for bioinformatic support; N. Wilson for assistance with the facilities and technical support; M. Rostovskaya for discussions around eutherian amnion. We would also like to thank Mark Keller (Gehlenborg lab) for help in setting up the Vitessce visualization platform. M.-L.N.T. is funded by a Herchel Smith PhD Fellowship in Science. D.K. is funded by the Wellcome Mathematical Genomics and Medicine Programme at the University of Cambridge (PFZH/158 RG92770). C.G. was funded by the Swedish Research Council (2017-06278, administered by Sahlgrenska Cancer Center, University of Gothenburg). Work in the Gottgens group is supported by Wellcome, Bloodwise, MRC, CRUK, by core support grants from Wellcome to the Wellcome-MRC Cambridge Stem Cell Institute. Work in the Marioni group is supported by core funding from CRUK (C9545/A29580) and by the European Molecular Biology Laboratory. This work was funded as part of a Wellcome grant (220379/B/20/Z) awarded to B.G, J.N.. and J.C.M. Work in the E.B.G. lab is supported by CRUK (C9545/A29580).

## Author Contributions

C.G., M.-L.N.T. and J.N., performed embryo dissections and generated the scRNA-seq atlas dataset. M.-L.N.T. and F.J.C. generated the scATAC-seq atlas dataset. J.A.-R. And T.K.A supplied the animals for the experiments and conducted histology and imaging of the embryos. M.-L.N.T performed pre-processing and initial low-level analysis for both atlases. D.K. performed batch correction and global visualization of the scRNA-seq dataset. D.K. performed cross-species comparisons between the rabbit mouse, macaque, and human datasets. D.K. implemented the website. B.T performed analysis on the scATAC-seq dataset. D.K., M.-L.N.T., B.P-S., C.G., B.T., and I.I., annotated atlas cell types. M-L.N.T., B.T., and D.K., analyzed the atlas trophoblast cells. J.N., E.B.-G., J.M., B.G, supervised the study. M.-L.N.T., D.K., B.T., B.G., J.M., and E.B.-G. wrote the manuscript. All authors read and approved the final manuscript.

## Ethics declarations

Provision of time-mated rabbit embryos by Labcorp Early Development Laboratories Limited as well as single cell RNA sequencing costs were supported by a research contract agreement with Novo Nordisk A/S.

