## Supplementary Figures for "Rabbit Development as a Model for Single Cell Comparative Genomics"

**This PDF file includes:**

Figs S1 to S9
Table S1

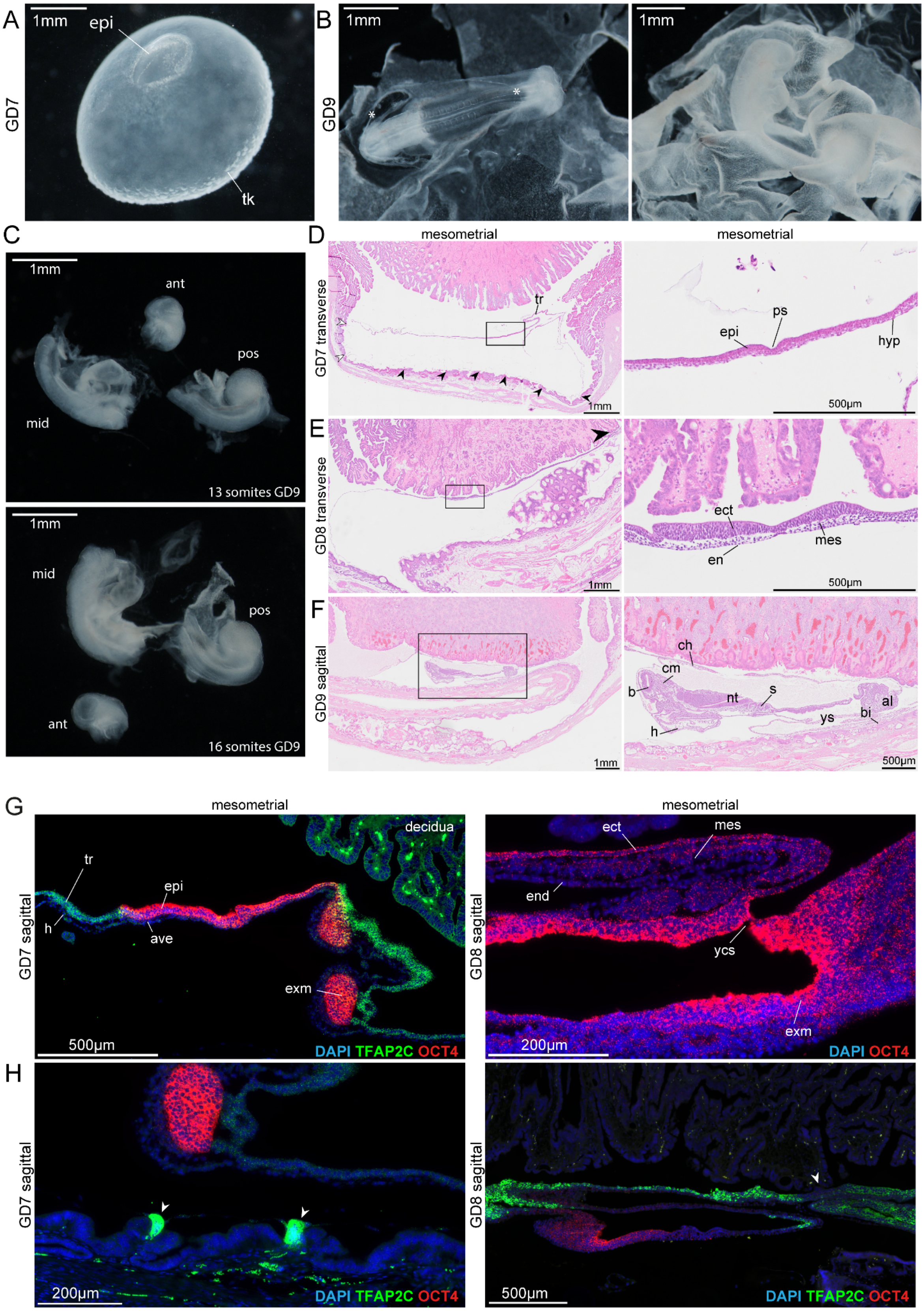
A
GD7
1mm
epi
tk
B
GD9
1mm
1mm
C
1mm
ant
pos
mid
13 somites GD9
1mm
mid
pos
ant
16 somites GD9
D
GD7 transverse
mesometrial
tr
1mm
mesometrial
ps
epi
hyp
500µm
E
GD8 transverse
1mm
ect
mes
en
500µm
F
GD9 sagittal
1mm
ch
cm
b
nt
s
al
bi
ys
h
500µm
G
GD7 sagittal
mesometrial
decidua
tr
epi
h
ave
exm
500µm
DAPI TFAP2C OCT4
GD8 sagittal
mesometrial
ect
mes
end
ycs
exm
200µm
DAPI OCT4
H
GD7 sagittal
200µm
DAPI TFAP2C OCT4
GD8 sagittal
500µm
DAPI TFAP2C OCT4

**Figure S1. Histology and dissection images reveals early dynamics of early germ layers of rabbit flat-disk embryo**

**A-C**, Brightfield images of rabbit dissection showing early structures in development. **A**, GD7 pre-implantation rabbit blastocyst, containing an exposed epiblast layer and visible primitive streak on one side trophoblast and trophoblastic knobs on the other side (See fig 1a-c). The blastocyst shows considerable inflation. **B**, GD9 rabbit embryos showing amniogenesis (left, asterisk), early organogenesis and yolk sac (right). right = rabbit embryo with visible optic vesicle, heart, and surrounding extraembryonic tissues. **C**, images of a 13 somite (top) and 16 somite (bottom) GD9 rabbit embryo microdissected into anterior, mid, and posterior regions. The anterior region contains the cranial section, the mid portion contains the heart and adjacent structures, and the posterior includes the allantois. **D-F**, Histology images of rabbit cross-sections in uterus for rabbit at GD7, GD8, and GD9 shown in Figure 1C. The mesometrial side is oriented towards the top of the page, while the anti-mesometrial side is oriented towards the bottom. **D**, left = zoomed out image showing early embryo layer with implantation sites in the anti-mesometrial side (arrows). right = zoomed in photo of the epiblast, underlying hypoblast, and primitive streak. tr, trophoblast; ps, primitive streak; hyp, hypoblast; epi, epiblast. **E**, transverse GD8 rabbit embryo section showing signs of early implantation on the mesometrial side (arrow, left). (right) Zoom of the embryo-proper with the three germ layers visible. ect, ectoderm; mes, mesoderm; end, endoderm. **F**, Sagittal section of a GD9 rabbit in uterus (left). Enlarged image (right) annotates various developing organs. h, heart; b, brain; cm, craniac mesoderm; ch, chorion; nt, neural tube; s, somites; ys, yolk sac; bi, blood island; al, allantois. **G**, RNAscope images of sagittal rabbit sections with DAPI, *TFAP2C* and *OCT4* (left) or DAPI and *OCT4* (right). *TFAP2C* is highly expressed in the trophoblast (left), the epiblast and extraembryonic mesoderm express *OCT4*, and the hypoblast and anterior visceral endoderm are negative for both *TFAP2C* and *OCT4*. ave= anterior visceral endoderm; exm= extraembryonic mesoderm; ycs= yolk connecting stalk. **H**, Trophoblast *OCT4* and *TFAP2C* expression in sagittal sections of a GD7 rabbit embryo. Trophoblastic knobs adhering to the uterine lining at mesometrial implantation sites are visible as *TFAP2C*+ clusters.The *OCT4*+ cluster of cells are extraembryonic mesoderm, as in F; left. Overlying chorioamnion trophoblast can be seen as *TFAP2C*+ cells (right). Additionally, some weakly *OCT4*+ cells can be seen which are cell types originating from the epiblast. Arrows indicate fusion between the fetal and maternal layers.

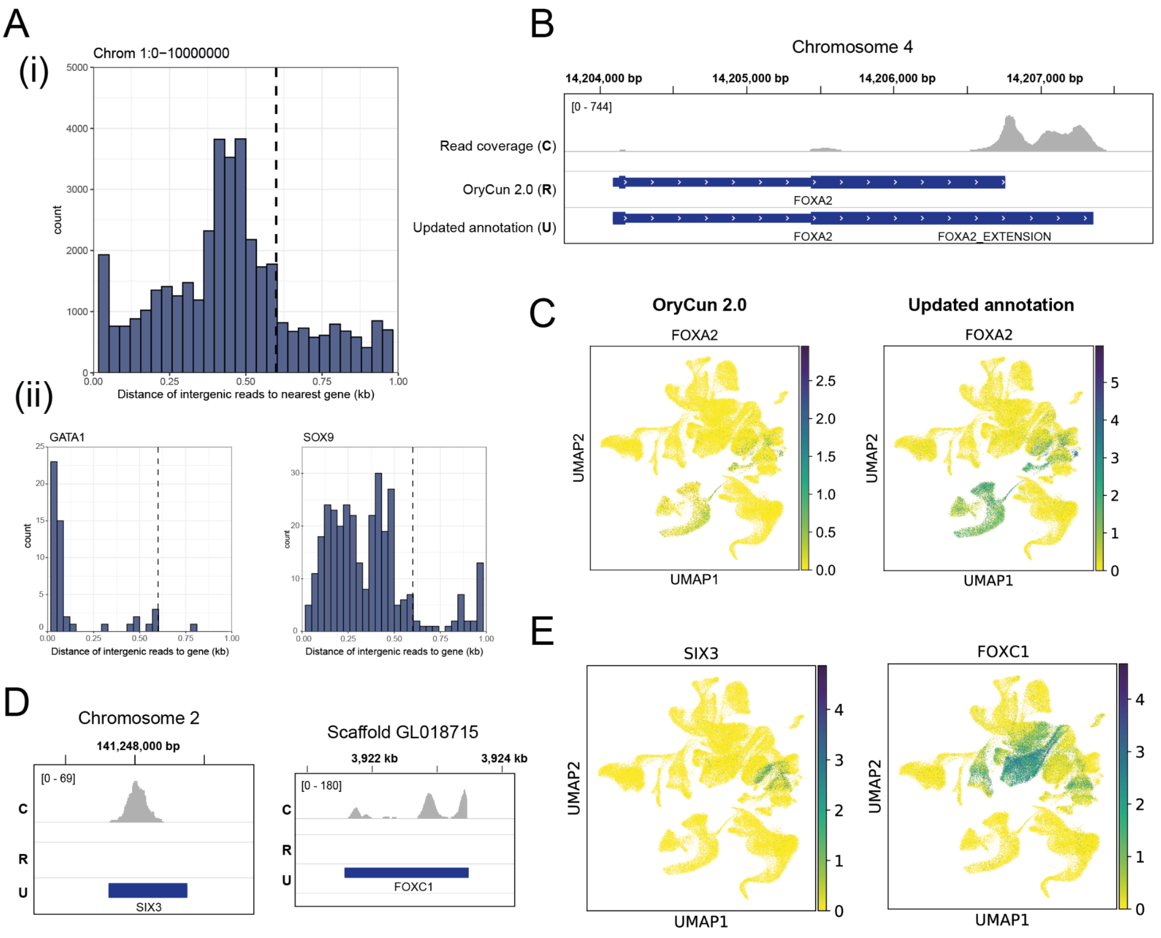


**Figure S2. Updated transcriptome annotation improves the quality of processed scRNA-seq data.**

**A**, The distances of intergenic reads to their nearest annotated gene in a single GD8 sample. **i**) The distances for a subset of intergenic reads aligned to a region of chromosome 1 are shown. The vertical line indicates the 600bp extension added to the 3' ends (see Methods). **ii**) Distances are also shown for intergenic reads whose closest annotated gene is *GATA1* and *SOX9*. **B**) The read coverage for the same GD8 sample shows that many reads associated with *FOXA2* transcripts are positioned off the 3' end of the OryCun 2.0 reference annotation. These missing reads are captured by our 600bp extension. **C**) The 3' extended gene annotations substantially improve the signal obtained after processing with cellranger. **D**) Examples of new annotations added from aligning human gene annotations with the rabbit transcriptome. These new annotations overlap positions with high read coverage. **E**) The signal captured as a result of the new human gene annotations are consistent with the known expression patterns of *SIX3* (forebrain marker) and *FOXC1* (mesodermal marker).

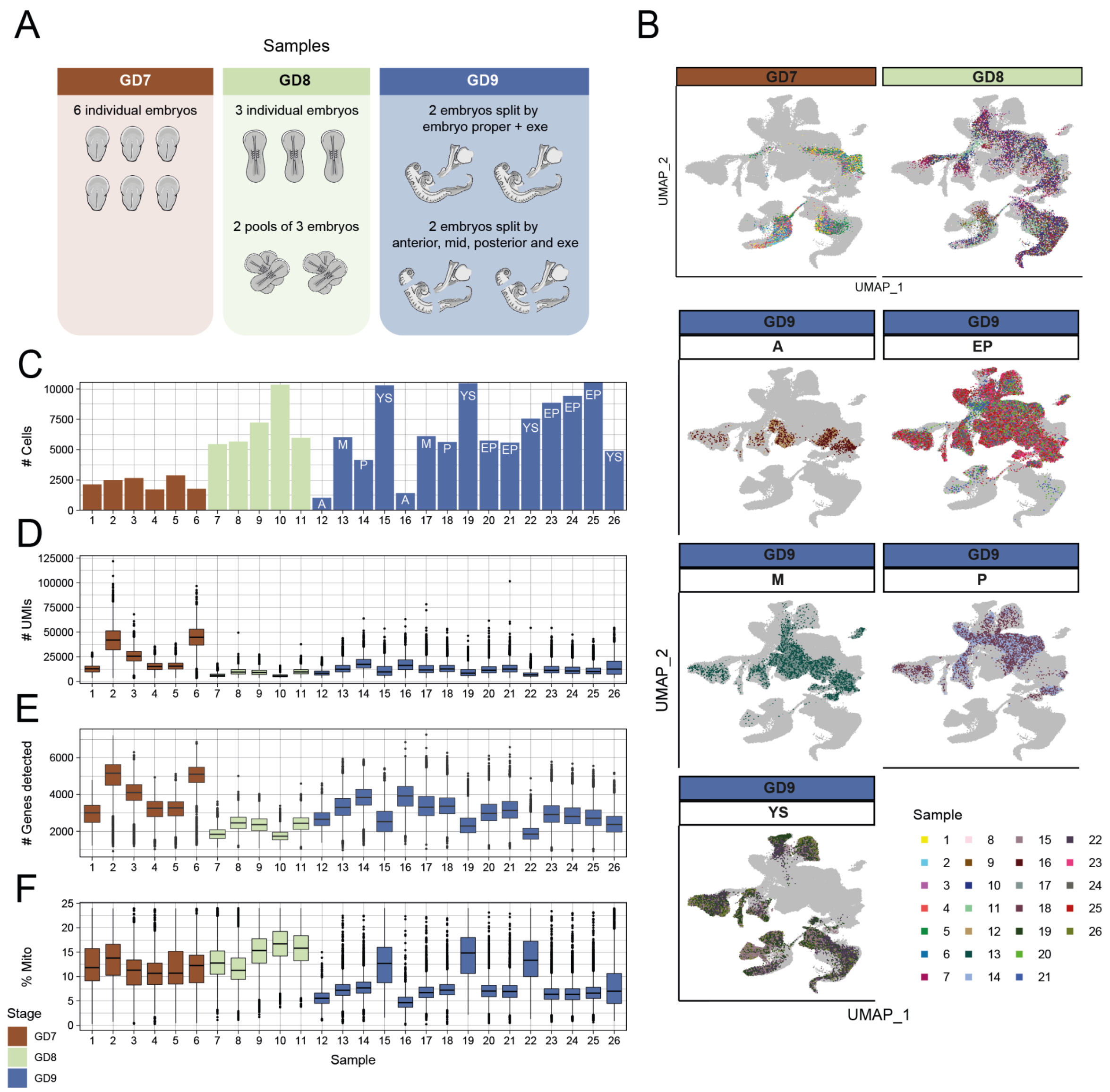


**Figure S3. RNA-seq quality controls.**

**A**, An illustration of the samples processed for single-cell RNA-sequencing. **B**) Cells from samples taken at each developmental stage and anatomical dissection are highlighted in the UMAP embedding. Following batch correction, the samples seem well mixed within the UMAP visualisation. Number of cells (**C**), UMIs (**D**), genes detected (**E**) and percentage of mitochondrial reads (**F**) across each of the samples, coloured by developmental stage. Annotations in **C** refer to the anatomical dissections performed for GD9 samples. A -

anterior section, M - mid section; P - posterior section; YS -yolk-sac/extraembryonic tissues; EP - embryo proper.

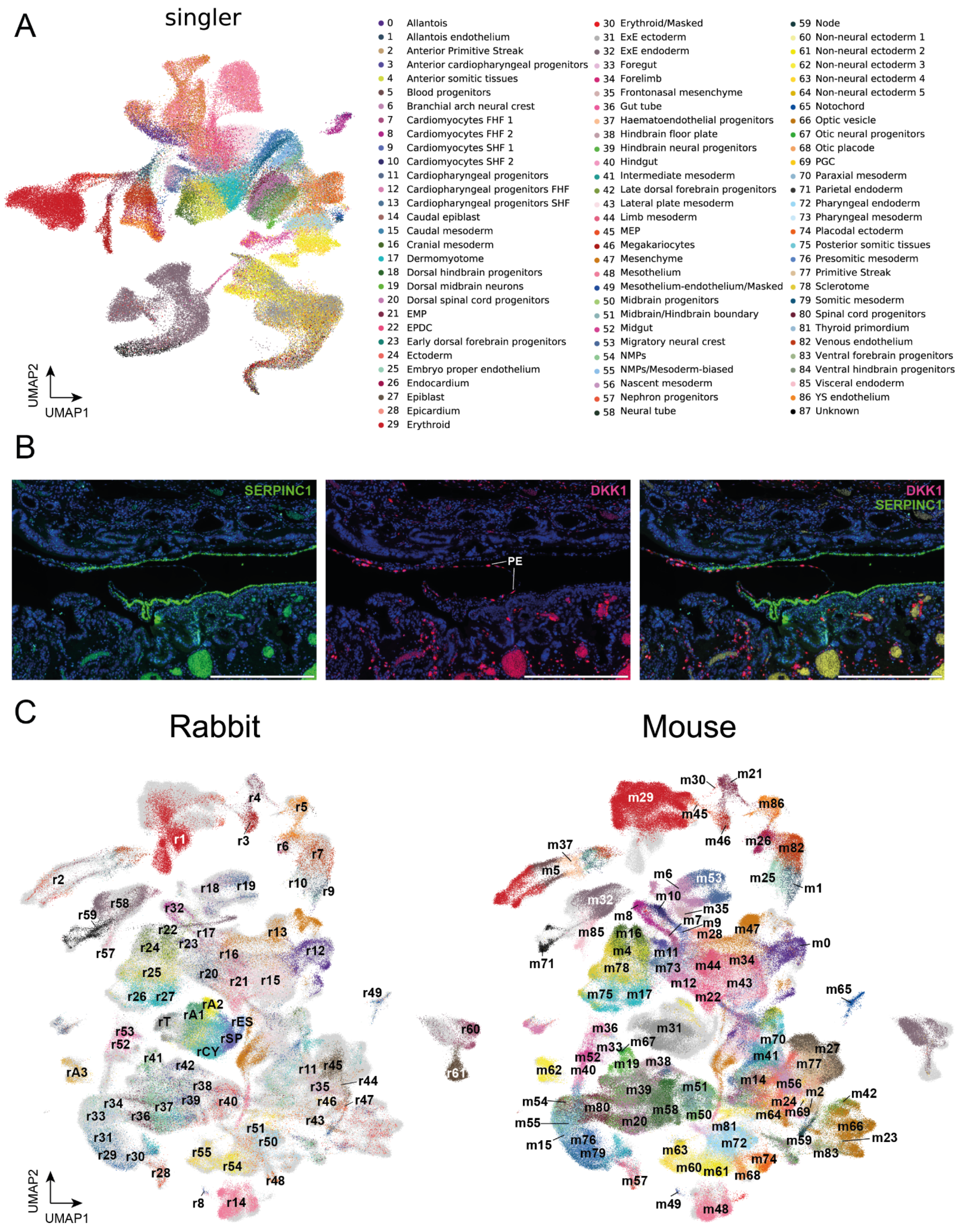


**Figure S4. Cell type annotation.**

**A**, SingleR predicted cell type annotations for each cell of the rabbit atlas, following training on the mouse atlas. Predicted labels are more easily visualised via the interactive web app, accessible via https://marionilab.github.io/RabbitGastrulation2022/. **B**) RNAscope images of *SERPINC1* and *DKK1* helped identify cells of the parietal endoderm (PE). Scale bars = 500 µm. **C**) Cells of the rabbit (left) and mouse (right) (Ross-Handler et al. 2022) datasets in a SAMap integrated UMAP embedding. Cells are coloured and annotated according to the cell type labels of each respective atlas. The rabbit (r) and mouse (m) label numberings refer to those in figure legends 1C and S4A respectively. Shortened labels for the rabbit extra-embryonic ectoderm cell types have also been added. rT = Trophoblast; rA1 = Amnion 1; rA2 = Amnion 2; rA3 = Amnion 3; rCY = Cytotrophoblast; rSP = SCT progenitors; rES = Early SCT.

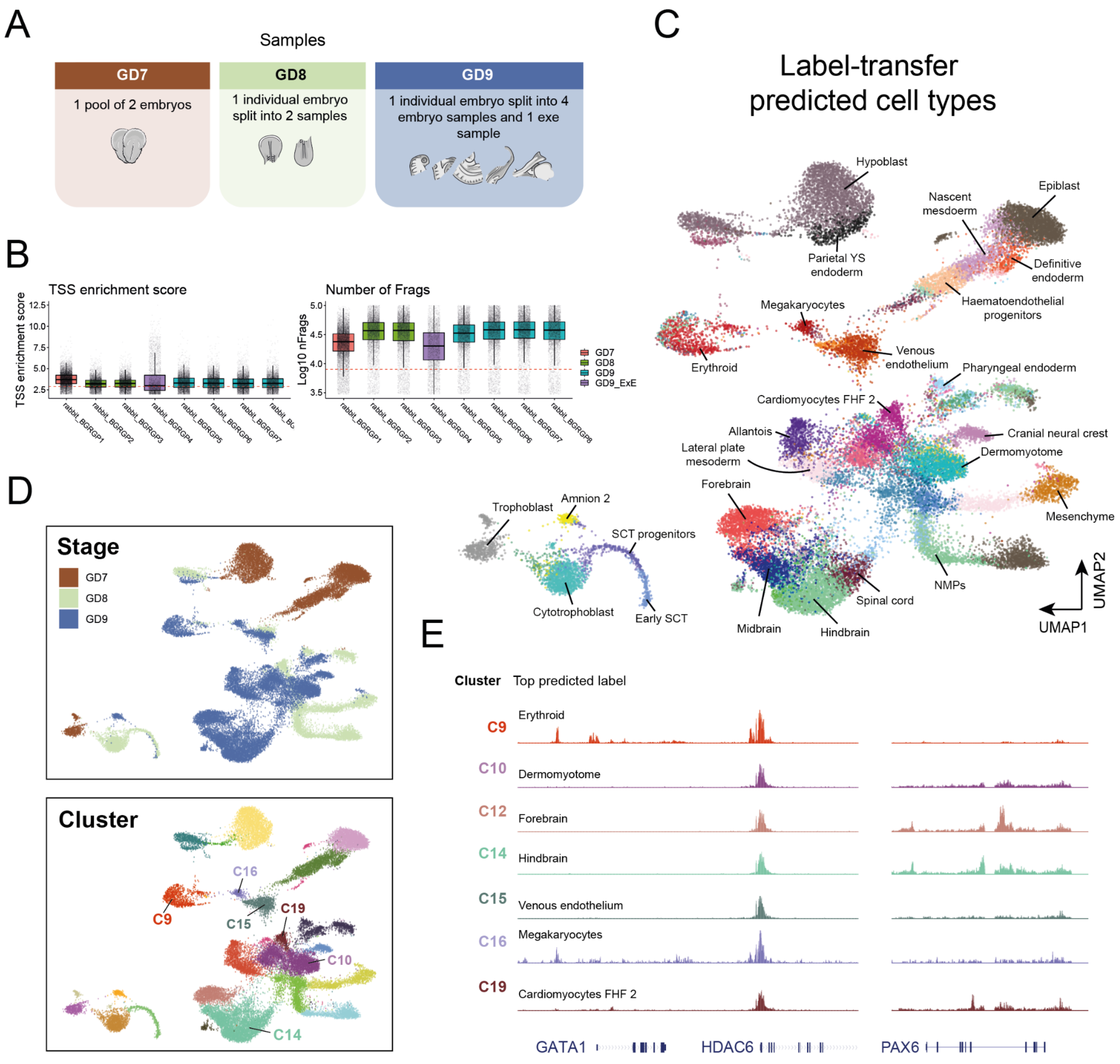


**Figure S5. Whole embryo scATAC-seq and trophoblast annotation.**

**A**) An illustration of the samples processed for single-cell ATAC-sequencing. **B**) Quality control of whole embryo scATAC-seq cells based on transcription start site (TSS) enrichment score and total number of fragments per cell. Red dotted lines indicate thresholds set for cells to be included in the analysis. **C**) UMAP of 34,082 cells that passed quality control for scATAC-seq, colored by predicted cell type inferred from label transfer (see Methods). **D**) UMAP from C colored by developmental stage (top), and cluster (bottom). **E**) Genome browser views of the regions surrounding GATA1, HDAC (left) and

PAX6 (right) for a subset of the identified clusters. The predominant cell type prediction across cells of each cluster is also shown which validates accessibility in blood and neural clusters.

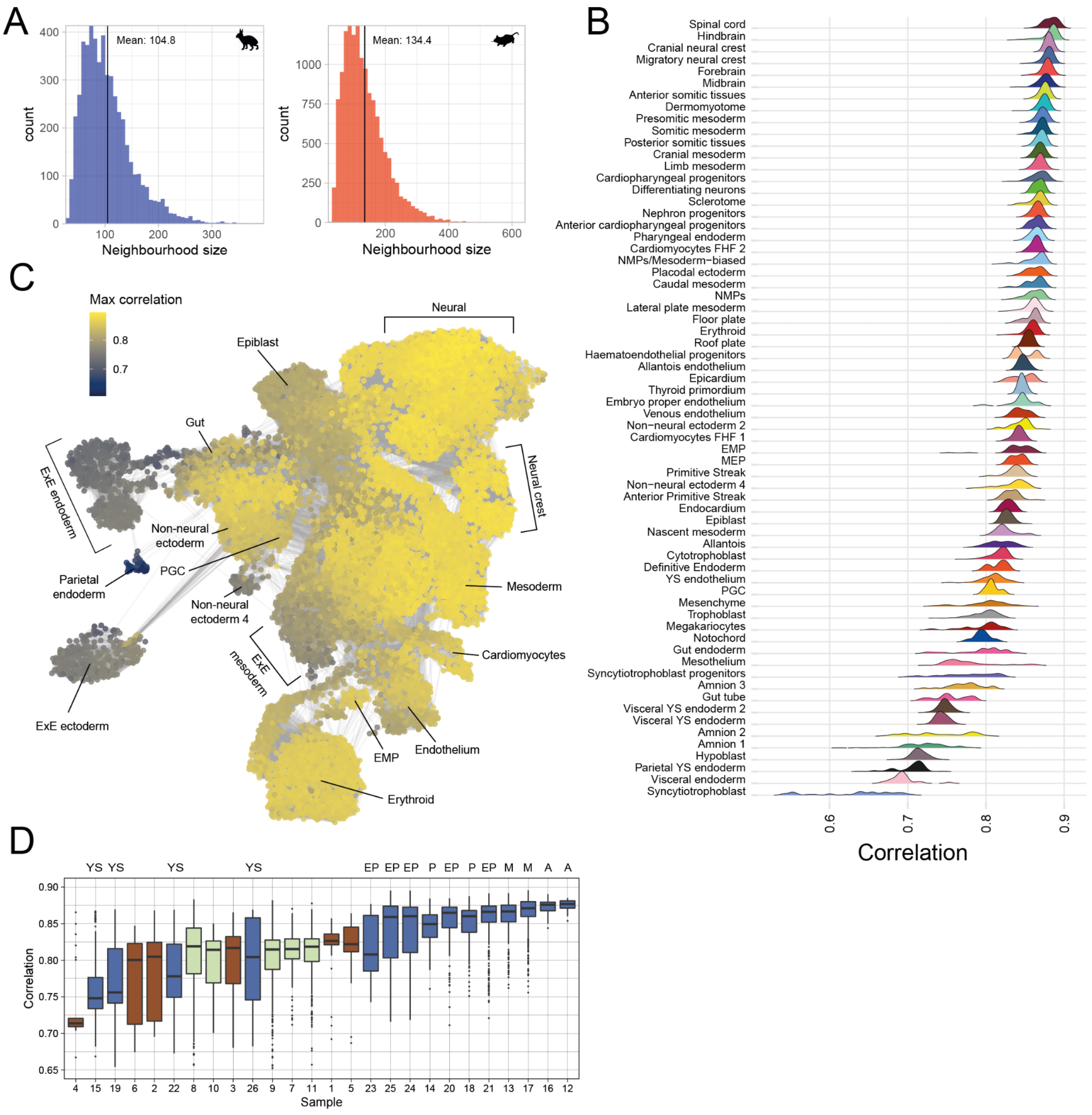


**Figure S6. Rabbit-mouse neighborhood comparisons.**

**A,** Distribution of the number of cells within neighbourhoods of the rabbit and mouse atlas. B) Same plot as in Figure 3D shown for all cell types. Extraembryonic tissues tend to trend lower on the similarity score with the highest similarity scoring cell types corresponding to spinal cord, hindbrain, and cranial neural crest. C) Neighbourhoods of the mouse atlas coloured according to their maximum correlation with any rabbit neighbourhood. General UMAP positions for a subset of cell types are shown.. D) Maximum correlation scores, grouped by the sample of each neighbourhood index cell. Samples are coloured by the developmental stage and annotations above the GD9 samples refer to the anatomical dissection. A - anterior section, M - mid section; P - posterior section; YS -yolk-sac/extraembryonic tissues; EP - embryo proper.

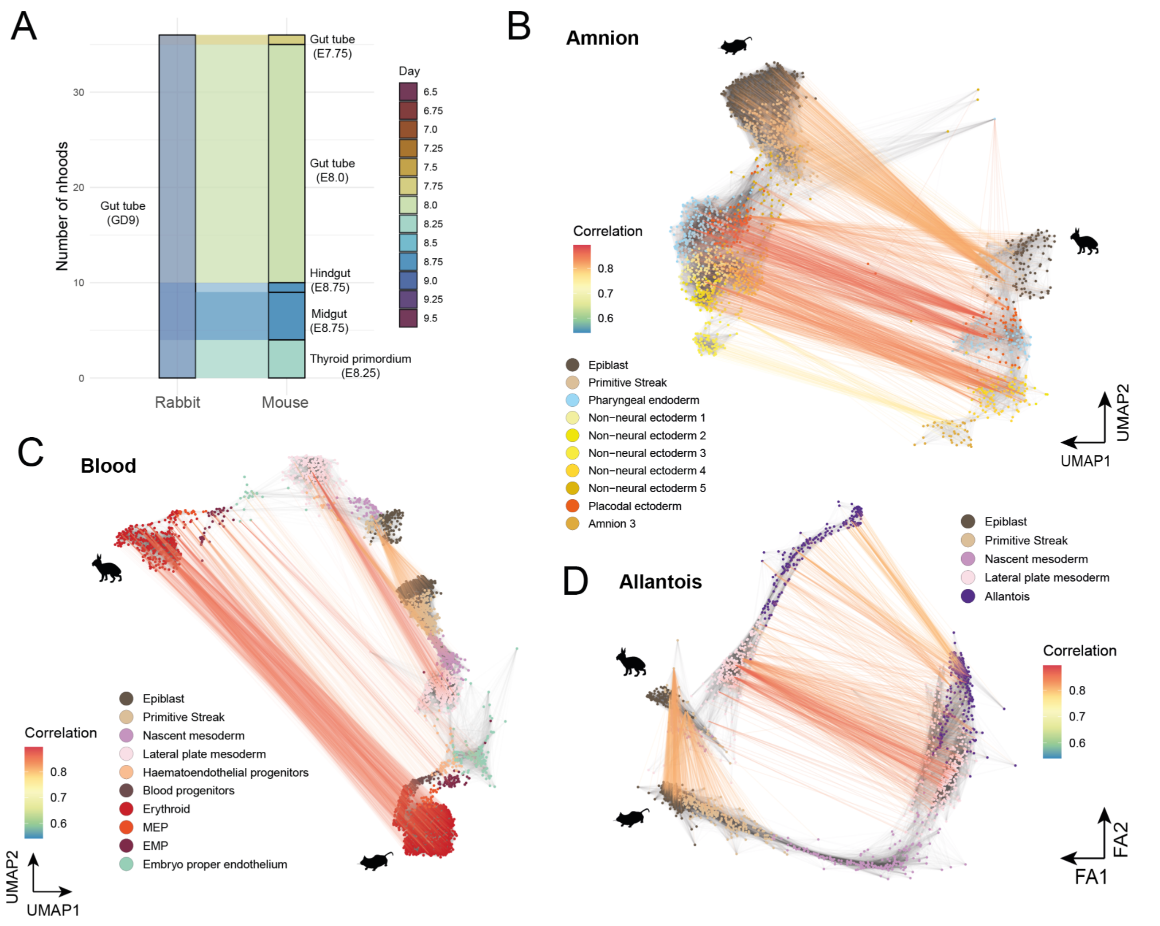


**Figure S7. Neighbourhood comparisons along differentiation trajectories.**

**A)** Alluvial plot showing the proportion of rabbit gut tube neighbourhoods which form maximally correlated mappings with mouse neighbourhoods of different cell types. **B-D)** Neighbourhoods associated with **B**) amnion, **C**) erythroid and **D**) allantois differentiation trajectories are shown for the rabbit and mouse. Maximally correlated neighbourhood pairs are linked, with the strength of correlation indicated with line colour. Neighbourhoods are coloured by annotated cell type. Neighborhoods in **B**,**C** are represented in the UMAP embedding, whereas in **D**, neighborhoods are positioned according to a ForceAtlas2 graph layout.

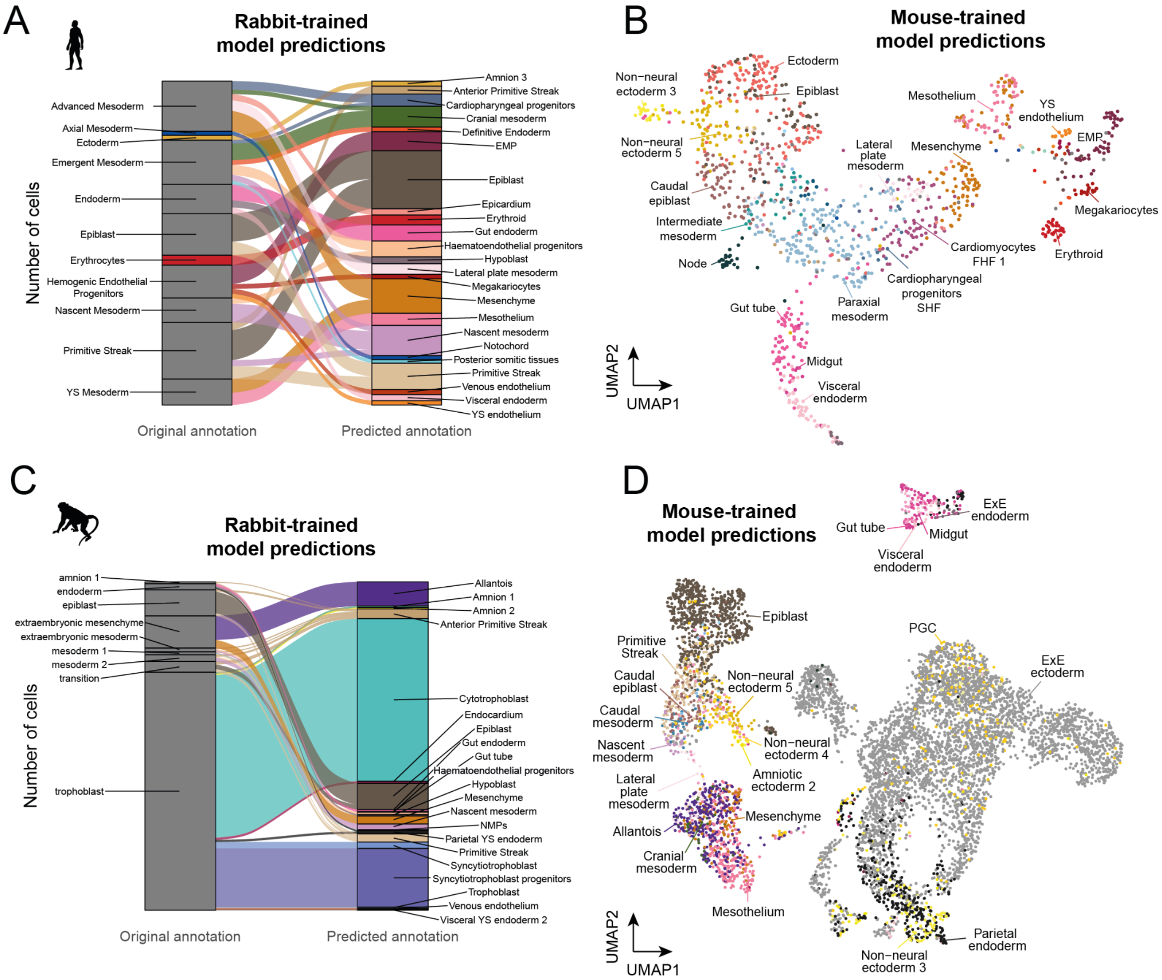


**Figure S8. Human and macaque cell type prediction with SingleR.**

**A**) Alluvial plot showing the changes to the original cell type labels after predicting annotations with a SingleR model trained on the rabbit atlas. Mappings only shown for predicted cell types with more than 10 cells. **B**) UMAP of the CS7 human dataset (as in Tyser et al. 2021, Figure 1C) coloured according to the SingleR cell type predictions trained on the mouse atlas. **C**) Same as in **A** for the classification of cells from the macaque in-vitro dataset (as in Yang et al. 2021, Figure 1b). **D**) Same as in **B** for the Yang et al. 2021 dataset.

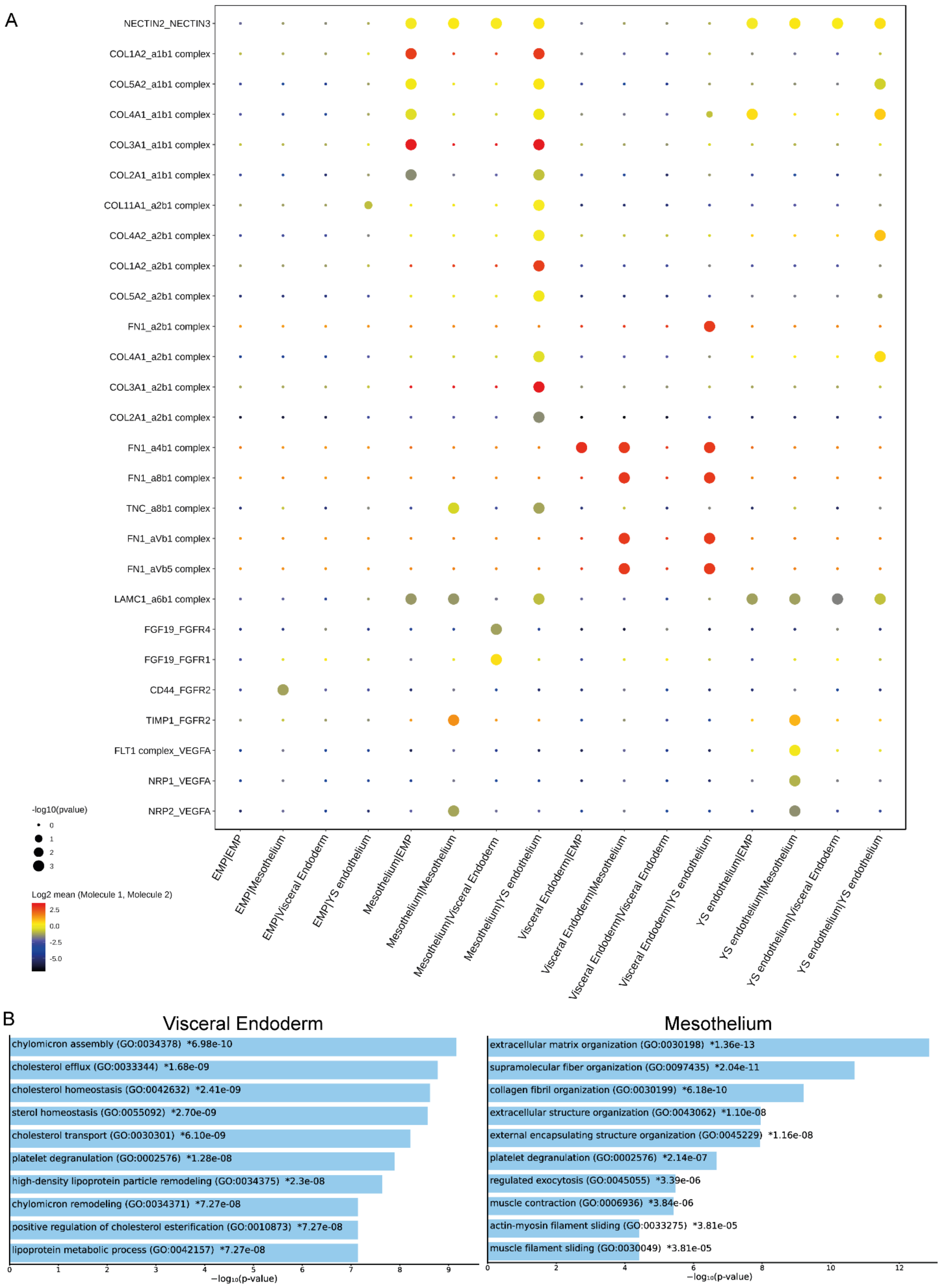
A
NECTIN2_NECTIN3
COL1A2_a1b1 complex
COL5A2_a1b1 complex
COL4A1_a1b1 complex
COL3A1_a1b1 complex
COL2A1_a1b1 complex
COL11A1_a2b1 complex
COL4A2_a2b1 complex
COL1A2_a2b1 complex
COL5A2_a2b1 complex
FN1_a2b1 complex
COL4A1_a2b1 complex
COL3A1_a2b1 complex
COL2A1_a2b1 complex
FN1_a4b1 complex
FN1_a8b1 complex
TNC_a8b1 complex
FN1_aVb1 complex
FN1_aVb5 complex
LAMC1_a6b1 complex
FGF19_FGFR4
FGF19_FGFR1
CD44_FGFR2
TIMP1_FGFR2
FLT1 complex_VEGFA
NRP1_VEGFA
NRP2_VEGFA
-log10(pvalue)
0
1
2
3
Log2 mean (Molecule 1, Molecule 2)
2.5
0.0
-2.5
-5.0
EMP|EMP
EMP|Mesothelium
EMP|Visceral Endoderm
EMP|YS endothelium
Mesothelium|EMP
Mesothelium|Mesothelium
Mesothelium|Visceral Endoderm
Mesothelium|YS endothelium
Visceral Endoderm|EMP
Visceral Endoderm|Mesothelium
Visceral Endoderm|Visceral Endoderm
Visceral Endoderm|YS endothelium
YS endothelium|EMP
YS endothelium|Mesothelium
YS endothelium|Visceral Endoderm
YS endothelium|YS endothelium
B
Visceral Endoderm
chylomicron assembly (GO:0034378) *6.98e-10
cholesterol efflux (GO:0033344) *1.68e-09
cholesterol homeostasis (GO:0042632) *2.41e-09
sterol homeostasis (GO:0055092) *2.70e-09
cholesterol transport (GO:0030301) *6.10e-09
platelet degranulation (GO:0002576) *1.28e-08
high-density lipoprotein particle remodeling (GO:0034375) *2.3e-08
chylomicron remodeling (GO:0034371) *7.27e-08
positive regulation of cholesterol esterification (GO:0010873) *7.27e-08
lipoprotein metabolic process (GO:0042157) *7.27e-08
0 1 2 3 4 5 6 7 8 9
−log10(p-value)
Mesothelium
extracellular matrix organization (GO:0030198) *1.36e-13
supramolecular fiber organization (GO:0097435) *2.04e-11
collagen fibril organization (GO:0030199) *6.18e-10
extracellular structure organization (GO:0043062) *1.10e-08
external encapsulating structure organization (GO:0045229) *1.16e-08
platelet degranulation (GO:0002576) *2.14e-07
regulated exocytosis (GO:0045055) *3.39e-06
muscle contraction (GO:0006936) *3.84e-06
actin-myosin filament sliding (GO:0033275) *3.81e-05
muscle filament sliding (GO:0030049) *3.81e-05
0 2 4 6 8 10 12
−log10(p-value)

**Figure S9. Ligand-receptor interactions and gene ontology reveal roles of visceral endoderm and mesothelium in yolk sac hematopoietic niche.**

**A**, Extended dot plot of CellphoneDB output reveals a large network of extracellular matrix protein interactions between the mesothelium and endothelium, as well as VEGF and FGF complexes. The order of the cell types along the x-axis indicates the order of the molecule1:molecule2 expression along the y axis. The size of the dot indicates the -$\log_{10}$(pvalue), while the color of the dot indicates the $\log_2$ mean of the mean expression of the molecules. Of note, YS endothelium and mesothelium show a significant interaction between FLT1 and FLT family complexes with VEGFA, which plays a known role in haematopoiesis. **B**, Gene ontology analysis of the differential transcriptional expression of the visceral endoderm clusters and the mesothelium clusters reveal different roles in the yolk sac niche. Visceral endoderm cells score highly on nutrient transport roles like chylomicron assembly, cholesterol efflux, and cholesterol homeostasis (left). Mesothelium transcripts interact highly with the endothelium according to (A), and score highly on extracellular matrix organization and collagen fibril organization. This indicates a closer interaction with endothelium that is critical for the function of the yolk sac hematopoietic niche and the mesothelium provides a role in structure separate from the visceral endoderm cells. P-values are adjusted for multiple-testing.

**Table S1: Table_S1.pdf (separate file)**

Cell numbers and quality statistics for each scRNA-seq and scATAC-seq sample.
