## Supplementary Table 1 for "Rabbit Development as a Model for Single Cell Comparative Genomics"

| Assay | 10X sample | Barcode | time-point | notes | somite number | total cell/nuclei counts in sample | estimated number of loaded cells/nuclei in 10X lane | estimated captured (60% of loaded) | cellranger called cells/nuclei | passed QC |
|---|---|---|---|---|---|---|---|---|---|---|
| scRNAseq | 1 | SIGAA9 | GD7 | GD7_embryo1 | 0 | 9768 | 6956 | 4174 | 4941 | 2139 |
| scRNAseq | 2 | SIGAB9 | GD7 | GD7_embryo2 | 0 | 24500 | 16450 | 9870 | 8197 | 2501 |
| scRNAseq | 3 | SIGAC9 | GD7 | GD7_embryo3 | 0 | 14300 | 6721 | 4033 | 5581 | 2661 |
| scRNAseq | 4 | SIGAD9 | GD7 | GD7_embryo4 | 0 | 5250 | 3525 | 2115 | 3215 | 1714 |
| scRNAseq | 5 | SIGAE9 | GD7 | GD7_embryo5 | 0 | 7150 | 5170 | 3102 | 5184 | 2885 |
| scRNAseq | 6 | SIGAF9 | GD7 | GD7_embryo6 | 0 | 22640 | 13301 | 7981 | 4935 | 1774 |
| scRNAseq | 7 | SIGAA11 | GD8 | GD8_embryo1 | 4 | 85000 | 12484 | 7491 | 14147 | 5456 |
| scRNAseq | 8 | SIGAB11 | GD8 | GD8_embryo2 | 4 | 108800 | 16495 | 9897 | 14270 | 5670 |
| scRNAseq | 9 | SIGAC11 | GD8 | GD8_pool1 (3 pooled embryos) | 0 | 116800 | 17708 | 10625 | 30,523 | 7237 |
| scRNAseq | 10 | SIGAD11 | GD8 | GD8_pool2 (3 pooled embryos) | 0 | 69550 | 11674 | 7005 | 21033 | 10342 |
| scRNAseq | 11 | SIGAE11 | GD8 | GD8_embryo3 | 4 | 99200 | 15040 | 9024 | 20059 | 5981 |
| scRNAseq | 12 | SIGAA12 | GD9 | GD9_embryo1_anterior | 13 | 2990 | 2008 | 1205 | 2642 | 1039 |
| scRNAseq | 13 | SIGAB12 | GD9 | GD9_embryo1_mid | 13 | 40375 | 15181 | 9109 | 13644 | 6024 |
| scRNAseq | 14 | SIGAC12 | GD9 | GD9_embryo1_posterior | 13 | 25900 | 8695 | 5217 | 12098 | 4142 |
| scRNAseq | 15 | SIGAD12 | GD9 | GD9_embryo1_ExE | 13 | 39600 | 15510 | 9306 | 24881 | 10295 |
| scRNAseq | 16 | SIGAE12 | GD9 | GD9_embryo2_anterior | 16 | 3080 | 2895 | 1737 | 3307 | 1420 |
| scRNAseq | 17 | SIGAF12 | GD9 | GD9_embryo2_mid | 16 | 26600 | 8930 | 5358 | 10204 | 6113 |
| scRNAseq | 18 | SIGAG12 | GD9 | GD9_embryo2_posterior | 16 | 19250 | 8225 | 4935 | 11088 | 5632 |
| scRNAseq | 19 | SIGAH12 | GD9 | GD9_embryo2_ExE | 16 | 54000 | 14100 | 8460 | 24482 | 10473 |
| scRNAseq | 20 | SIGAE8 | GD9 | GD9_embryo3_embryo_proper | 11 | 30000 | 11750 | 7050 | 3676 | 5744 |
| scRNAseq | 21 | SIGAF8 | GD9 | GD9_embryo3_embryo_proper | 11 | 30000 | 11750 | 7050 | 9469 | 5576 |
| scRNAseq | 22 | SIGAG8 | GD9 | GD9_embryo3_ExE | 11 | 31800 | 14946 | 8968 | 14923 | 7557 |
| scRNAseq | 23 | SIGAH8 | GD9 | GD9_embryo4_embryo_proper | 19 | 105300 | 17675 | 10605 | 19,114 | 8861 |
| scRNAseq | 24 | SIGAF11 | GD9 | GD9_embryo4_embryo_proper | 19 | 105300 | 17675 | 10605 | 19,306 | 9425 |
| scRNAseq | 25 | SIGAG11 | GD9 | GD9_embryo4_embryo_proper | 19 | 105300 | 17675 | 10605 | 21255 | 10578 |
| scRNAseq | 26 | SIGAH11 | GD9 | GD9_embryo4_ExE | 19 | 22800 | 8930 | 5358 | 12,013 | 4894 |
| scATACseq | BGRGP1 | SINAA3 | GD7 | 2 GD7 embryos pooled | 0 | 46870 | 16000 | 9600 | 10593 | 8750 |
| scATACseq | BGRGP2 | SINAB3 | GD8 | GD8_embryo1 | 4 | 160437 | 16000 | 9600 | 5675 | 4082 |
| scATACseq | BGRGP3 | SINAC3 | GD8 | GD8_embryo1 | 4 | 160437 | 16000 | 9600 | 6025 | 4336 |
| scATACseq | BGRGP4 | SINAD3 | GD9 | Extraembryonic | 18 | 35625 | 16000 | 9600 | 3453 | 1629 |
| scATACseq | BGRGP5 | SINAE3 | GD9 | GD9_embryo1 | 18 | 280125 | 16000 | 9600 | 4742 | 4064 |
| scATACseq | BGRGP6 | SINAF3 | GD9 | GD9_embryo1 | 18 | 280125 | 16000 | 9600 | 4784 | 3994 |
| scATACseq | BGRGP7 | SINAG3 | GD9 | GD9_embryo1 | 18 | 280125 | 16000 | 9600 | 4553 | 3771 |
| scATACseq | BGRGP8 | SINAH3 | GD9 | GD9_embryo1 | 18 | 280125 | 16000 | 9600 | 4074 | 3456 |
